# Stress-responsive *Mycobacterium tuberculosis* subpopulations manipulate macrophage polarization and can be targeted to limit inflammation

**DOI:** 10.64898/2026.03.13.711673

**Authors:** Laura Pokorny, Lalit Kumar Singh, Nicolas Gangneux, Margaux Pillon, Claudia Chica, Natalia Pietrosemoli, Emeline Perthame, José Crispin Zavala Alvarado, Olivier Hebert, Christophe Rochais, Cédric Lecoutey, Alban Lepailleur, Mena Cimino, Etienne Kornobis, Marvin Albert, Jean-Yves Tinevez, Eléonore Bouscasse, Mariette Matondo, Stéphane Lo, Zélie Julia, Sylvie Le Gac, Nathalie Grall, Giulia Manina

## Abstract

Tuberculosis is characterized by broad clinical heterogeneity that hinders infection control, with differences in lesion development, progression, and treatment outcomes. This complexity is likely associated with *Mycobacterium tuberculosis* inherent phenotypic variation and its capacity to diversify under host microenvironmental and antimicrobial stressors. Here, we analyze *M. tuberculosis* at the single-cell and subpopulation level using fluorescent reporters, imaging, transcriptomic, and functional assays. We identify RNA signatures specific to stress-responsive bacilli with translational potential. Focusing on the clinically validated chaperone GroEL2, we find that it correlates with *M. tuberculosis* growth rate and stress tolerance in vitro and intracellularly. Furthermore, GroEL2 phenotypic diversity influences innate responses in macrophages, which experience different polarization, in turn affecting GroEL2 expression. We also show that targeting GroEL2 impairs pathogen survival and dampens inflammation. This study provides a link between pathogen phenotypic variation and macrophage fates, with implications for early infection outcomes, local disease progression, and subpopulation-targeted interventions.

## Introduction

Tuberculosis remains a major public health threat, ranking among the top ten causes of mortality and being the leading cause of death from a single infectious agent (Warner et al., 2025; Sati et al., 2025). Although its prevalence is highest in low- and middle-income countries, the transmission of both drug-susceptible and increasingly drug-resistant *Mycobacterium tuberculosis* continues worldwide. Treatment is particularly lengthy due to the inherent ability of *M. tuberculosis* to endure stressful conditions and the emergence of drug resistance, leading to severe side effects and lower success rates (Dheda et al., 2024; Dartois et al., 2025). Thus, rifampicin-resistant *M. tuberculosis* is recognized as a critical priority pathogen by the World Health Organization, considerably contributing to the antimicrobial resistance (AMR) crisis, and responsible for approximately one in five AMR-related deaths (Sati et al., 2025). Tackling this global health challenge entails better understanding of the complexity of tuberculosis across multiple scales and leveraging this knowledge to guide interventions (Warner et al., 2025).

Infection with *M. tuberculosis* can follow distinct clinical trajectories, ranging from early pathogen clearance to a continuum of asymptomatic forms, which may eventually reactivate, and symptomatic forms, which primarily affect the lungs but can disseminate to other organs (Simmons et al., 2018; Scriba et al., 2024). This clinical heterogeneity occurs not only among patients but also within the same individual, as separate lung lesions can evolve and respond to therapy independently, making it difficult to predict disease severity, risk of progression, and treatment response (Flynn and Chan, 2022). Indeed, spatial analyses of pulmonary lesions revealed marked cellular, molecular, transcriptional, immunological, and inflammatory heterogeneity, as well as uneven drug distribution, implying that local host microenvironments influence infection dynamics both between and within lesions (Marakalala et al., 2016; Lavin and Tan, 2022; Fonseca et al., 2025). These divergent local responses are likely to arise during the initial stages of infection, when tissue-resident and recruited macrophages adopt distinct polarization states dependent on ontogeny, spatial context, immune stimuli, and the pathogen’s traits, thus influencing early bacterial control and downstream immune activation (Ehrt et al., 2018; Bain and MacDonald, 2022; Russell et al., 2025). Single-cell studies have revealed that, despite its population clonality (Gagneux et al., 2018), *M. tuberculosis* unevenly expresses various cellular traits under optimal growth conditions, and that this cell-to-cell variation increases under environmental perturbations (Dhar et al., 2016; Manina et al., 2019; Mishra et al., 2021; Chung et al., 2022; Sherry and Rego, 2024). A hallmark of *M. tuberculosis* physiology is the variation in replication and metabolic state both in vitro and in vivo (Manina et al., 2015; Ehrt et al., 2018; Mishra and Saito, 2022). The capacity of genetically identical bacilli to dynamically expand their phenotypic repertoire increases their ability to withstand microenvironmental challenges, with consequences for bacterial persistence, disease progression, and therapeutic efficacy (Mekonnen et al., 2021; Lavin and Tan, 2022; Rutschmann et al., 2022).

A major hindrance to the control of tuberculosis is that standard molecular, culture-based, and immunological diagnostics (Kontsevaya et al., 2024) fail not only to fully capture the spectrum of tuberculosis infection but also to identify functionally distinct *M. tuberculosis* phenotypic variants. Although phenotypic heterogeneity is recognized as central to *M. tuberculosis* physiology and fitness (Dhar et al., 2016; Mishra et al., 2021; Chung et al., 2022; Sherry and Rego, 2024), its relationship with divergent clinical trajectories remains unclear. Greater understanding of this phenomenon at the bacterial level and in the context of infection will help clarify mechanisms critical to the survival of the pathogen and conceive more effective disease control strategies.

In vivo, *M. tuberculosis* encounters several conditions that disrupt the redox homeostasis of the cell, and adaptation to these perturbations is crucial to its survival (Mishra et al., 2021; Parbhoo et al., 2022). Ferredoxins are iron–sulfur (Fe-S) cluster proteins that mediate redox reactions, alleviate damage caused by redox stress, and support cellular homeostasis (Buckel and Thauer, 2018). One of them, FdxA (Rv2007c), controlled by the dormancy regulon (Schnappinger et al., 2003; Kendall et al., 2004), is a small ferredoxin that contains two Fe-S clusters and mediates electron transfer at low redox potential (Ricagno et al., 2007). Even though FdxA is not essential in exponentially growing cultures, it was reported to be strongly upregulated under stressful conditions, such as hypoxia or low pH, and during late stages of infection, making it a suitable marker of adaptation (Schnappinger et al., 2003; Rohde et al., 2007; Kesavan et al., 2009).

In this work, we show that clonal *M. tuberculosis* subpopulations expressing higher FdxA levels exhibit distinct transcriptional signatures consistent with increased stress responsiveness. Among the most upregulated signatures in these subpopulations, we identify GroEL2, an essential heat-shock protein involved in protein homeostasis and host immunomodulation (Sia and Rengarajan, 2019). Furthermore, we find that GroEL2 is upregulated in the sputum of patients with subclinical tuberculosis and during treatment, supporting its relevance for *M. tuberculosis* persistence. Surprisingly, *M. tuberculosis* cells expressing higher GroEL2 levels grow faster both in vitro and intracellularly and have a fitness advantage under host-mimetic stressors and anti-tubercular drugs. Using single-cell and subpopulation analyses, we show that pre-existing variation in GroEL2 levels predicts differential macrophage polarization states, which in turn influence GroEL2 expression. Additionally, bacilli engineered to produce high levels of GroEL2 oligomers elicit strong pro-inflammatory responses from different cellular compartments, involving Toll-like receptor (TLR) 4. Finally, we show that targeting GroEL2 decreases host cell inflammation without impairing pathogen inhibition by standard antibiotic treatment. Our study proves that dynamic variation in *M. tuberculosis* GroEL2 expression is associated not only with differential stress responsiveness but also with distinct innate responses. These findings have implications for local development of infection and bacterial persistence and point to GroEL2 as a relevant biomarker at the host–pathogen interface.

## Results

### Stress-responsive *M. tuberculosis* subpopulations exhibit distinct transcriptomic signatures

Host microenvironments expose *M. tuberculosis* to fluctuations in nutrient availability, oxygen levels, and antimicrobial molecules, disrupting its metabolic and redox homeostasis (Wilburn et al., 2018; Pacl et al., 2018). Survival under these conditions requires extensive physiological adaptations, including increased levels of Fe-S clusters, which are associated with drug tolerance (Mishra et al., 2021). Since FdxA is upregulated under several stress conditions in vitro and in vivo (Schnappinger et al., 2003; Rohde et al., 2007; Kesavan et al., 2009), we used it as a starting proxy for *M. tuberculosis* adaptation. To monitor FdxA expression at the single-cell level, we fused a red-fluorescent marker in frame with the FdxA-encoding gene under its native regulatory region, preserving the N-terminal Fe-S cluster-binding core (Ricagno et al., 2007). The resulting FdxA_mCherry_ reporter strain grew similarly to the control strain in axenic conditions (Figure S1A), showed high cell-to-cell fluorescence variation during exponential growth, and experienced further changes under host-mimetic conditions (Figure S1B).

To probe whether the inherent or stress-induced FdxA_mCherry_ variation was indicative of broader transcriptional reprogramming, we carried out low-input RNA sequencing (RNA-Seq) on subpopulations of bacilli sorted by FdxA_mCherry_ levels (Figures 1A and S1C). We enriched for FdxA_mCherry_-Dim or -Bright cells during exponential and stationary growth phases and after exposure to fatty acids as a sole carbon source or hypoxia, simulating conditions relevant to tubercular lesions (Wilburn et al., 2018; Pacl et al., 2018). Principal component (PC) analysis revealed that the main source of variance in our datasets is due to differences in culture conditions, as expected. Interestingly, PC also distinguished subpopulations within each culture condition, separating them into discrete clusters (Figure 1B). By comparing FdxA_mCherry_-Bright versus -Dim subpopulations (Table S1), we identified differentially expressed genes (DEGs) in all conditions (Figures 1C, 1D, and S1D), with the highest numbers observed under host-mimetic conditions, particularly propionate, a precursor of virulence lipids associated with host persistence and drug tolerance (Wilburn et al., 2018). While exposure to propionate and hypoxia resulted in the largest number of shared DEGs, regardless of the direction of expression (Figures 1C and 1D), subpopulation-specific transcriptional profiles in the exponential phase of growth and under propionate exposure showed the strongest positive correlation (Figure 1E). In contrast, subpopulations emerging during the stationary phase of growth displayed either no or negative correlation relative to other conditions, leading us to exclude it from further analysis (Figure 1E).

**Figure 1.**
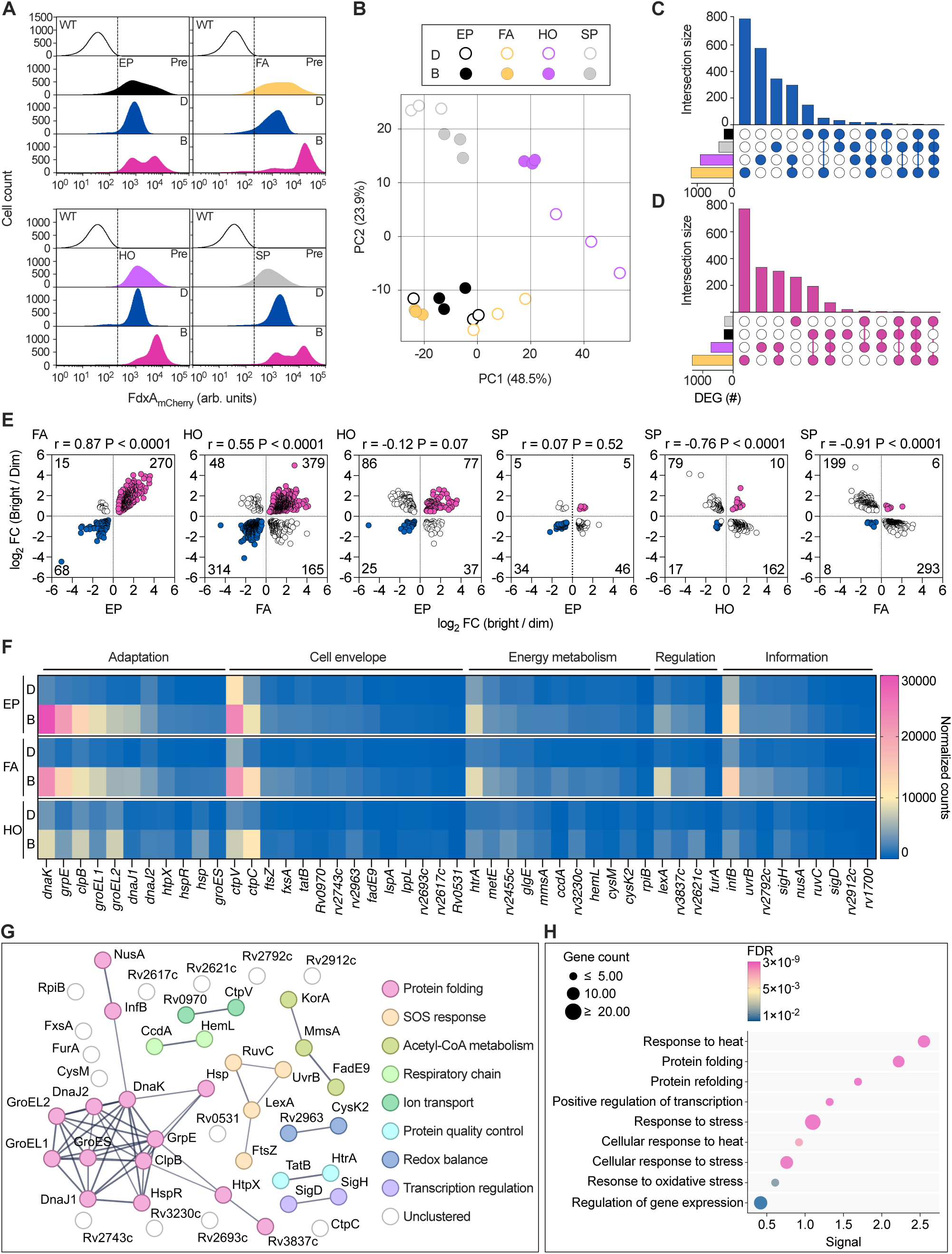
Sorted FdxA_mCherry_ clonal subpopulations exhibit different transcriptomic signatures **(A)** Representative flow cytometry density curves of wild-type (WT) *M. tuberculosis* (black line) and FdxA_mCherry_ reporter before (Pre) and after sorting of Dim (D) and Bright (B) subpopulations, in exponential phase (EP, black), fatty acid (FA, yellow), hypoxia (HO, lavender), and stationary phase (SP; gray) are normalized to 100,000 events (n = 3 independent experiments). **(B)** Principal component analysis (PCA) projection (n = 3) with percentages of variance associated with each axis indicated in brackets. FdxA_mCherry_-Dim (D, open circles) and -Bright (B, filled circles) subpopulations under conditions as in (A). **(C and D)** Upset plot of upregulated DEGs lists (filled circles) in each comparison, for FdxA_mCherry_-Dim (C) and -Bright (D) subpopulations. Vertical bars show the number of significant DEGs (FDR < 0.05, without fold-change threshold) specific to or shared between conditions color-coded as in (A and B). Horizontal bars show the total number of significant DEGs for each comparison. (E) Gene expression changes in the FdxA_mCherry_-Bright versus -Dim subpopulation for all significant DEGs (P < 0.05) between different condition pairs. DEGs per quadrant, Pearson correlation, and corresponding P values are indicated. (F) Average of normalized counts for genes significantly upregulated in FdxA_mCherry_-Bright (B) versus -Dim (D) subpopulations, shared among the indicated conditions. DEGs are clustered into functional categories (top line) and ranked by expression level in the EP-B condition. (G) STRING protein interaction network (PPI enrichment P < 1.0^e–16^) of genes significantly upregulated in the FdxA_mCherry_-Bright subpopulation (log_2_ fold-change (FC) ≥ 1, FDR < 0.05) and shared across EP, FA, and HO conditions (https://version-12-0.string-db.org/cgi/network?networkId=bcn3ndTz5deS). Edges represent high-confidence functional associations (interaction score ≥ 0.7), and line thickness indicates confidence strength. Nodes are grouped by k-means clustering into color-coded clusters. (H) Gene Ontology enrichment analysis of biological processes for the network in (G). Terms are grouped by similarity (≥ 0.8) and sorted by enrichment strength (≥ 0.01) with minimum gene count of 2. Circle size reflects the number of genes per term, and color gradient indicates FDR (≤ 0.05). Enriched processes highlight functional pathways associated with the FdxA_mCherry_-Bright’s gene set.

According to our initial assumption, we focused on genes that were significantly upregulated in FdxA_mCherry_-Bright subpopulations, which were expected to be more stress responsive and prone to adaptation than dim ones. Forty-nine differentially expressed genes (DEGs) were shared among the three remaining conditions and belonged to five main functional classes (Figure 1F and Table S2). Protein interaction and gene ontology analyses indicated enrichment in pathways related to transcriptional regulation, acetyl-CoA metabolism and respiration, metal ion transport, redox balance, stress responses, and protein quality control (Figures 1G, 1H, and S1E). These results show that clonal *M. tuberculosis* cells comprise phenotypically diverse subpopulations, which exhibit specific transcriptional signatures both prior to and following stress exposure, and that transcriptional remodeling associated with higher FdxA levels reflects a state of stress responsiveness that may support adaptation.

### *M. tuberculosis* stress-response biomarkers in sputum correlate with patient disease status

Sputum is widely used as a proxy for the lower respiratory tract to detect *M. tuberculosis* through phenotypic and genotypic bacteriological analyses, despite technical hindrances (Meehan et al., 2019). To explore the translational potential of *M. tuberculosis* subpopulation-specific signatures of stress responsiveness, we carried out a pilot observational study in a small cohort of patients with clinical tuberculosis, subclinical infection (defined as a positive IFN-γ release assay in the absence of clinical symptoms), and those undergoing either intensive- or continuation-phase treatment. To directly detect biomarker expression ex vivo, we analyzed RNA samples isolated from sputum remnants, collected during routine patient care, using the nCounter technology, without amplification steps. We designed 130 reporter probes targeting genes strongly upregulated in FdxA_mCherry_-Bright subpopulations (Figure 1).

However, because the fraction of biomarkers detected above the threshold varied across specimens, less than half of the patients originally recruited could be analyzed, i.e., four with clinical tuberculosis, two with subclinical infection, and four during different stages of treatment (Figure S2A), reducing the statistical power of the study. This was mainly due to the variable amount of sputum collected and handling time before freezing and to poor RNA yield and quality, typical of sputum samples. Moreover, matching specimens prior to and during treatment were available only in one case, limiting our ability to track intra-patient dynamics. Despite these drawbacks, PC analysis of gene expression data showed that patient groups explained about 40% of variance, while biomarker expression accounted for about 45% of total dataset variation (Figure 2A and Table S3). Unsupervised clustering stratified clinical cases versus subclinical and treated individuals, except for patient T#6, who had received treatment for less than one week at the time of sampling (Figure S2B). These results are consistent with the assumption that bacilli experience specific transcriptional reprogramming depending on disease severity and exposure to therapy.

**Figure 2.**
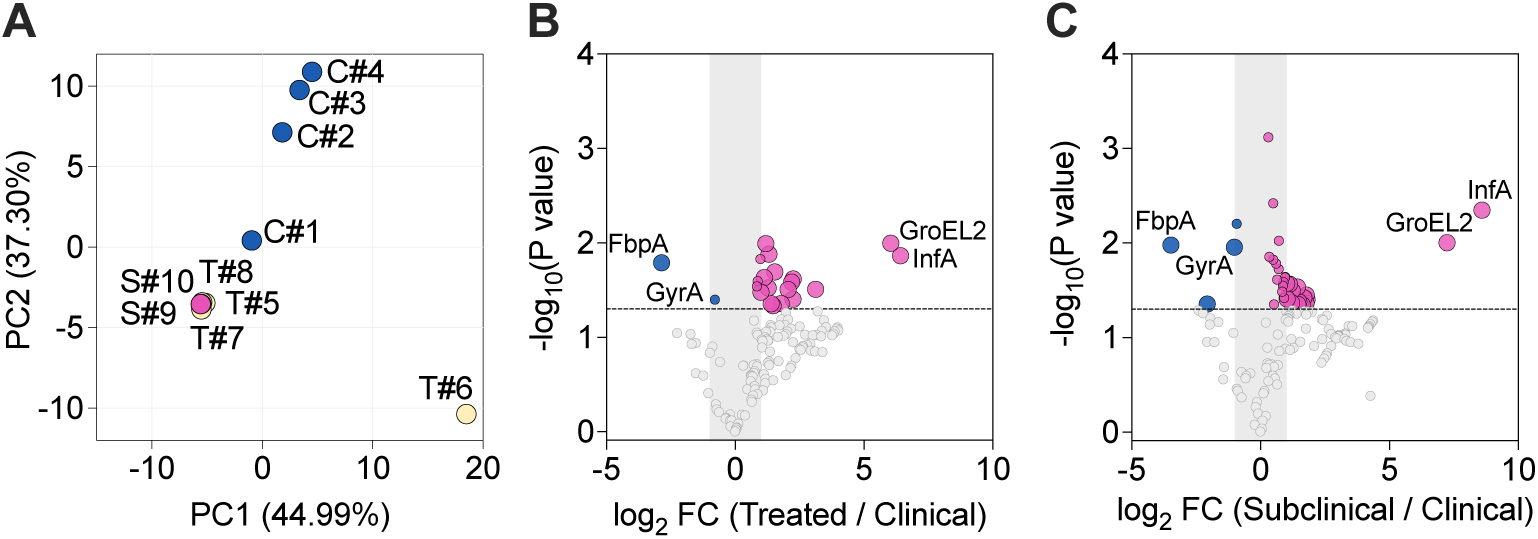
Analysis of *M. tuberculosis* stress-response biomarkers in patients’ sputum (A) PCA projection of *M. tuberculosis* gene expression data in sputum from ten patients with clinical (C, blue), subclinical (S, pink), or treated (T, yellow) tuberculosis. Normalized data were scaled to a mean of 0 and SD of 1. The first two PCs were selected, explaining 75% of total variance. Percentages of variance associated with each axis are indicated in brackets. **(B and C)** Volcano plots showing differential gene expression analysis comparing four treated (B) and two subclinical (C) with four clinical tuberculosis patients. Heteroscedastic Welch’s *t*-test was applied to log_2_-transformed counts (95% confidence interval). DEGs were ranked by -log_10_ of uncorrected P values and log_2_ FC. Dashed lines indicate P < 0.05 threshold, and shading log_2_ FC < 1.

Differential analysis identified sixteen biomarkers significantly upregulated at least twofold in treated versus clinical patients (Figure 2B) and fifteen in subclinical versus clinical patients (Figure 2C). Interestingly, five biomarkers, namely, *infA*, *groEL2*, *htrA, mutA*, and *rv3074*, were shared between both comparisons. Among them, *groEL2*, encoding the chaperonin CPN60-2, and *infA*, encoding the translation initiation factor 1, were the most strongly induced, suggesting a potential role in the emergence or maintenance of *M. tuberculosis* persistent subpopulations in vivo. The remaining shared biomarkers encode a putative serine protease involved in protein quality control (HtrA), a key enzyme in propionate metabolism (MutA), and an HNH nuclease-domain protein of unknown function (Rv3074). In summary, these results indicate that subpopulation-specific *M. tuberculosis* signatures of stress responsiveness can be detected in patients’ sputum and may be predictive of disease status. Despite the limitations of the study, including the small cohort size and suboptimal sample quality, we identified a subset of biomarkers upregulated in patients with subclinical tuberculosis and during therapy, suggestive of *M. tuberculosis* adaptations related to protein homeostasis, lipid metabolism, and DNA modification.

### Stress conditions alter *M. tuberculosis* GroEL2 expression, oligomerization, and localization

The subpopulation-specific transcriptional signatures of stress responsiveness identified in vitro included several genes encoding molecular chaperones; notably, GroEL2 was also upregulated in sputum from treated and subclinical patients (Figures 1 and 2). Chaperones are ubiquitous across all domains of life and maintain proteostasis by promoting folding, assembly, and translocation of proteins and protein complexes; additionally, they contribute to immunomodulation (Henderson et al., 2006). GroEL2, also known as heat shock protein 65 kDa (HSP65) or antigen A, is an essential and immunodominant chaperonin (Qamra et al., 2005). Structural studies indicate that *M. tuberculosis* GroEL2 is unlikely to assemble into tetradecameric oligomers typical of model organisms but instead primarily exists as dimers or cleaved monomers (Qamra et al., 2005; Shahar et al., 2011). Each monomer consists of an apical domain implicated in protein and host-cell binding, an intermediate hinge domain, and an equatorial domain mainly responsible for oligomerization and ATP binding (Figure S3A), albeit with limited ATPase activity (Qamra et al., 2005). GroEL2 was proposed to associate with the mycobacterial cell wall and to be secreted in either full-length or cleaved forms, with differential immunomodulatory roles (Sia and Rengarajan, 2019). Based on the relevance of GroEL2 in mycobacterial physiology and pathogenesis, we investigated its expression and cellular distribution in *M. tuberculosis*, using a C-terminal fluorescent reporter (GroEL2_mRuby_).

Single-cell microscopy of GroEL2_mRuby_ revealed heterogeneous patterns (Figures 3A–C). During exponential growth, GroEL2_mRuby_ localized as discrete fluorescence foci, predominantly at the cell poles. Upon exposure to subinhibitory concentrations of drugs targeting the cell wall or the cell cycle, or to hydrogen peroxide, GroEL2_mRuby_ showed either foci or a patchy distribution along the cell envelope with a significant increase in cell-to-cell heterogeneity and a decrease in mean fluorescence intensity (Figures 3B and S3B). Conversely, exposure to the transcription inhibitor rifampicin (RIF), the iron chelator deferoxamine (DFO), or nitric oxide (NO) caused a relocalization of GroEL2_mRuby_ along the cell length, with a strong increase under RIF and DFO and a decrease under NO (Figures 3C and S3B).

**Figure 3.**
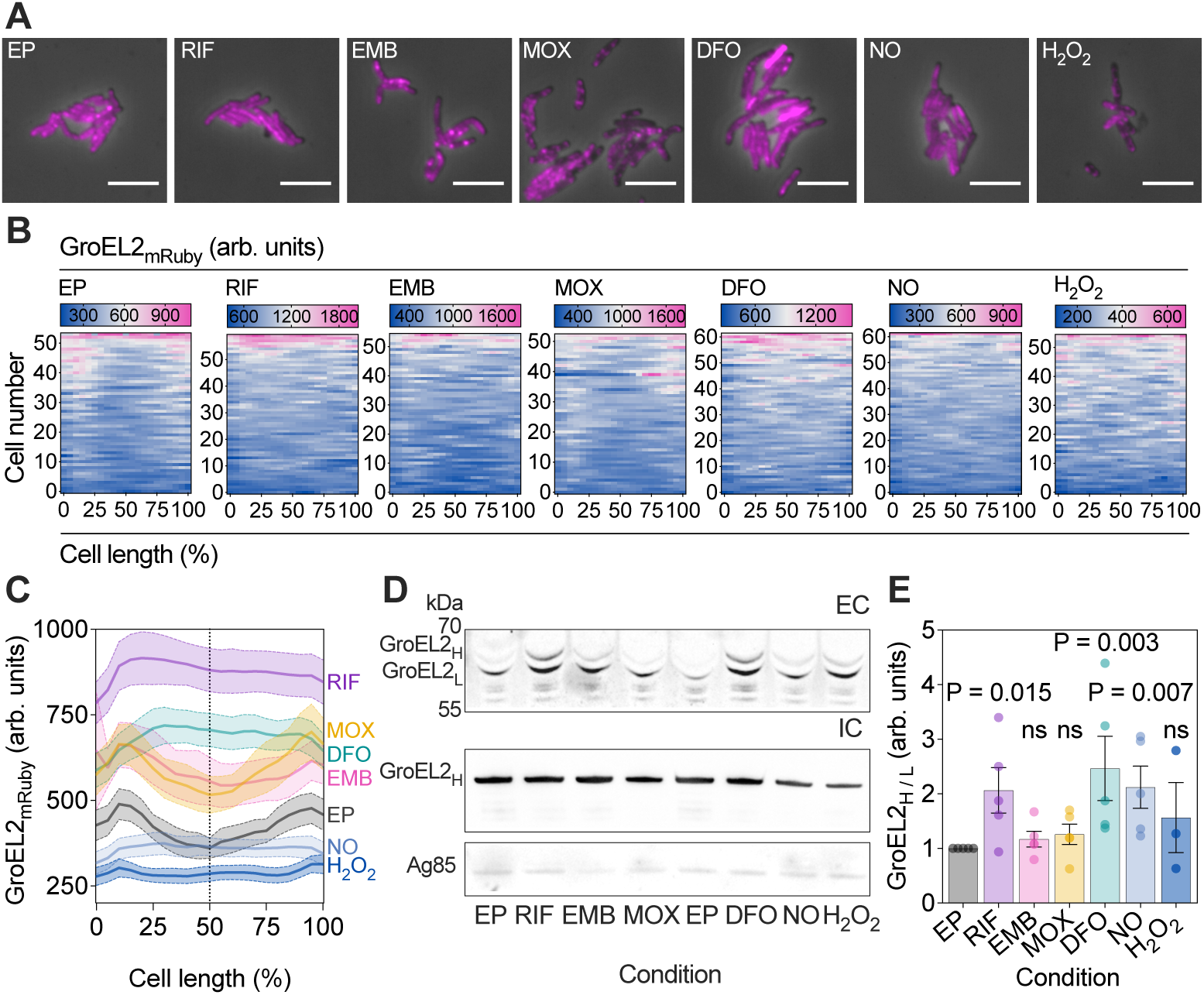
Stressful conditions alter GroEL2 expression, oligomerization, and localization (A) Representative snapshot images of *M. tuberculosis* GroEL2_mRuby_ reporter during exponential phase (EP) and after 48-hour exposure to subinhibitory concentrations of rifampicin (RIF, 0.038 µg/mL), ethambutol (EMB, 1.25 µg/mL), moxifloxacin (MOX, 0.025 µg/mL), deferoxamine (DFO, 0.25 mg/mL), nitric oxide (NO, 0.1 mM), or hydrogen peroxide (H_2_O_2_, 2.5 mM). Phase contrast (gray) and GroEL2_mRuby_ (magenta) are merged. Scale bars, 5 μm. (B) Heat maps showing GroEL2_mRuby_ intensity along the total cell length (as a percentage) for individual cells under the same conditions as in (A). Data are from two independent experiments. (C) Averaged single-cell traces of GroEL2_mRuby_ fluorescence along the percentage of total cell length (54 ≤ n ≤ 62 per condition). Population means and 95% confidence intervals (solid and dashed lines). (D) Western blot analysis of GroEL2 in wild-type *M. tuberculosis* grown under the same conditions as in (A). GroEL2 variants were detected by anti-HSP65 immunoblotting in extracellular (EC, supernatant) and intracellular (IC, cell lysate) fractions. EC fractions were normalized to bacterial cell density, and Ag85 was used as a loading control. The blot is representative of at least three independent experiments. (E) Quantification of the GroEL2_H/L_ in the EC fraction. Bars show mean ± SEM (3 ≤ n ≤ 5). Statistical significance by non-parametric Kruskal-Wallis test (P = 0.027), followed by multiple comparisons with two-stage linear step-up procedure of Benjamini, Krieger and Yekutieli (individual P values).

Next, we asked whether these distinct intracellular patterns were linked to differences in GroEL2 processing and release, since it was previously reported that the cell-envelope-associated serine hydrolase important for pathogenesis Hip1 (or CaeA), encoded by *rv2224c*, cleaves the full-length oligomeric form of GroEL2 into monomers at the cell envelope (Naffin-Olivos et al., 2014). Indeed, western blot analysis of intracellular and extracellular *M. tuberculosis* fractions revealed two main protein variants in the supernatant, namely, a higher-molecular-weight band (GroEL2_H_), consistent with the full-length oligomeric form, and a lower-molecular-weight band (GroEL2_L_), consistent with a cleaved monomeric form (Figure 3D). By contrast, the intracellular fraction contained exclusively the full-length oligomeric GroEL2_H_. Interestingly, RIF, DFO, and NO caused a significant increase in the extracellular GroEL2_H_ to GroEL2_L_ ratio (Figure 3E).

These observations support the hypothesis that intracellular foci may serve as both functional hubs and storage sites for GroEL2, which can be released extracellularly, with cleavage occurring or not depending on exogenous stimuli, including antibiotics and host-derived stressors. Although we could not determine the exact oligomeric state of the exported species, previous biochemical studies proposed that GroEL2 dimers expose free apical domains that may support both chaperone activity and interactions with host cells (Shahar et al., 2011). Overall, our findings support a model in which GroEL2 foci act as dynamic reservoirs from which GroEL2 can be mobilized under distinct forms, possibly endowing *M. tuberculosis* with different adaptive potential.

### Inherent GroEL2 heterogeneity predicts differential fitness in *M. tuberculosis* subpopulations

Because GroEL2 expression varies in response to different stressors in vitro and ex vivo (Figures 1-3), we asked whether inherent GroEL2 phenotypic variation is functional and contributes to *M. tuberculosis* adaptation to environmental fluctuations.

Flow cytometry analysis of exponentially growing GroEL2_mRuby_ reporter cells revealed two subpopulations, expressing either lower (Dim) or higher (Bright) fluorescence levels (Figures 4A and S4A), which further varied upon exogenous stimuli (Figure S4B). Surprisingly, time-lapse microscopy analysis showed relatively stable GroEL2_mRuby_ expression during the cell’s lifetime in exponential growth, typical of housekeeping genes (Manina et al., 2015), and a positive correlation between GroEL2_mRuby_ fluorescence and single-cell growth rate (Video S1, Figures 4B and 4C). Indeed, GroEL2_mRuby_-Bright cells (∼ 55%) grew significantly faster than -Dim ones (∼ 45%) (Figure 4D).

**Figure 4.**
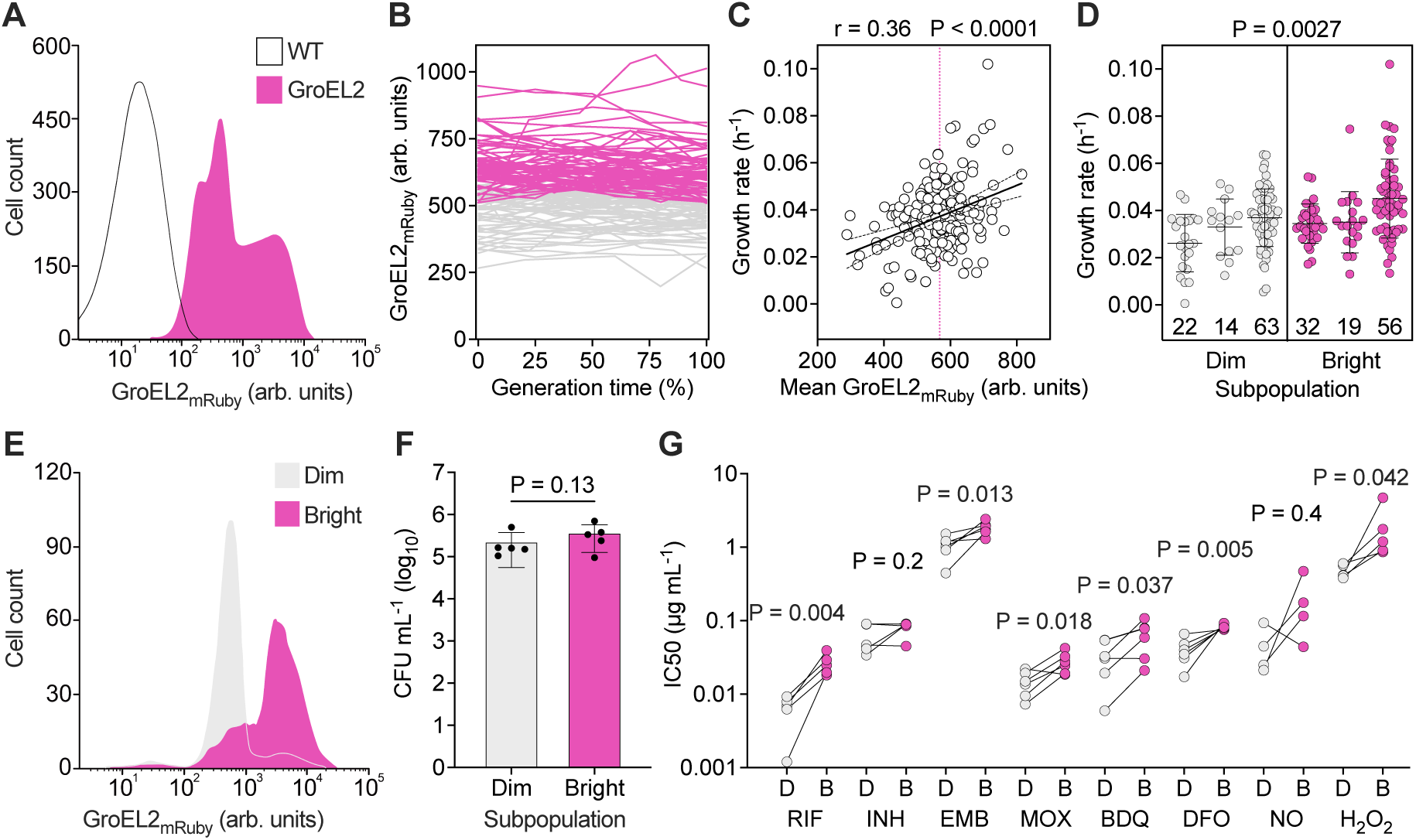
Inherent GroEL2_mRuby_ phenotypic variation predicts different *M. tuberculosis* fitness **(A)** Density curves of flow cytometry analysis of WT *M. tuberculosis* and GroEL2_mRuby_ reporter in exponential phase. Representative overlays are normalized to 100,000 events (n = 5). (B) Single-cell time traces of GroEL2_mRuby_ fluorescence expressed as a percentage of the generation time, separated into Dim (gray) and Bright (pink) based on average fluorescence (n = 2, with 148 individual cells). (C) Pearson’s correlation of single-cell growth rate with GroEL2_mRuby_ averaged over the cell’s lifetime (n = 3, with 206 individual cells). Population mean (dotted line). Simple linear regression fit (solid line, R^2^ = 0.14, P < 0.0001) and 95% confidence bands (dashed lines). (D) Single-cell growth rate of the 50% GroEL2_mRuby_ dimmest versus brightest cells. Lines represent mean ± SD (n = 3, with the number of cells shown at the bottom of the graphs). Significance by two-way ANOVA. (E) Density curves of flow cytometry analysis of GroEL2_mRuby_ fluorescence after subpopulation sorting. Representative overlays are normalized to 10,000 events (n = 5). (F) Cell density of sorted GroEL2_mRuby_ subpopulations. Lines represent mean ± SD (n = 5). Statistical significance by non-parametric two-tailed Wilcoxon signed rank test. (G) Half maximal inhibitory concentrations (IC50) of RIF, isoniazid (INH), EMB, MOX, bedaquiline (BDQ), DFO, NO, and H_2_O_2_ in flow-sorted GroEL2_mRuby_ Dim (D) and Bright (B) subpopulations. Lines represent mean ± SD of at least four independent experiments. Statistical significance by two-tailed paired *t*-test.

Next, we sorted exponentially growing GroEL2_mRuby_ cells by fluorescence, resulting in high post-sorting purity and comparable cell densities (Figures 4E, 4F, and S4C), and exposed these subpopulations to different anti-tubercular drugs or host-relevant stressors (Figures S4D and S4E). GroEL2_mRuby_-Bright cells showed enhanced survival under most conditions tested (Figure 4G). In summary, these results indicate that pre-existing variation in GroEL2 provides differential functional potential to *M. tuberculosis* subpopulations, with higher GroEL2 levels associated with both faster growth and increased tolerance to clinically relevant drugs and host-mimetic stresses.

## Intracellular GroEL2_mRuby_ phenotypic variation predicts distinct macrophage responses

Early recognition of pathogen-associated molecular patterns (PAMP) by pattern recognition receptors (PRRs) on phagocytes triggers innate immune responses, which influence inflammation and adaptive immunity, thus contributing to the fate of infection (Fitzgerald and Kagan, 2020; Daher et al., 2023). Molecular chaperones are potent immunogens and immunomodulators across kingdoms, although their phylogenetic distance ensures different mechanisms of immune recognition (Stewart and Young, 2004; Henderson et al., 2006). In *M. tuberculosis*, GroEL2 functions both as a protective chaperone and as a PAMP (Qamra et al., 2005; Sia and Rengarajan, 2019). Because differential GroEL2_mRuby_ expression correlated with distinct bacterial subpopulation fates (Figure 4), we asked whether this variation could also influence intracellular events during macrophage infection.

We infected naïve THP-1 macrophages (M0) with our GroEL2_mRuby_ reporter, which also constitutively expressed a fluorescent cytosolic (GFP_cyt_) marker (Figures 5A-5D and S5A-S5E). GFP_cyt_ allowed us to identify all infected cells, irrespective of GroEL2_mRuby_ levels, and to correct for technical sources of variation. Compared to exponentially growing bacilli, the GroEL2_mRuby_/GFP_cyt_ intensity ratio became more homogenous at 24 hours post-infection (hpi) and significantly decreased by 48 hpi (Figure 5E), suggesting dynamic regulation of this chaperone within the host cell. However, time-lapse imaging of infected macrophages (Video S2, Figures S5F and S5G) revealed relative fluorescence stability of single infectious foci over time (Figure 5F), allowing us to classify them as dimmer (ratio < 1) or brighter (ratio > 1). Consistent with GroEL2_mRuby_ single-cell growth kinetics (Figure 4C), the GroEL2_mRuby_/GFP_cyt_ ratio positively correlated with intracellular growth (Figure 5G), implying that higher GroEL2 levels provide a growth advantage to the pathogen inside the host cell. To examine the functional consequences of GroEL2 heterogeneity at the host–pathogen interface, we infected M0 macrophages for 24 hours and sorted them into GroEL2_mRuby_-Dim and - Bright subpopulations (Figures 5H and S5H-S5K), achieving high post-sorting purity (Figure S5L). Macrophages carrying GroEL2_mRuby_-Bright bacilli contained a relatively higher bacterial burden than their dim counterpart (Figure 5I), consistent with the faster growth rate of *M. tuberculosis* in both axenic culture and intracellularly (Figures 4C and 5G). Additionally, cytokine profiling at 24 hpi revealed moderate but significant polarization differences at the subpopulation level; namely, macrophages carrying GroEL2_mRuby_-Dim bacilli produced significantly more IL-10 and IL-4 than those carrying GroEL2_mRuby_-Bright bacilli, which produced significantly more TNF-α, but not IL-1β and IL-6 (Figure 5J). This suggested that macrophages infected with GroEL2_mRuby_-Dim bacilli tend toward a more permissive anti-inflammatory state, whereas those infected with GroEL2_mRuby_-Bright bacilli tend toward a more restrictive pro-inflammatory state. To further assess the influence of the macrophage activation status on GroEL2_mRuby_-GFP_cyt_ bacilli, we pre-polarized macrophages into either the M1 or M2 state prior to infection. As expected, pro-inflammatory M1 macrophages restricted the *M. tuberculosis* burden more efficiently than anti-inflammatory M2 macrophages (Figures S5M-S5O), but GroEL2_mRuby_ levels were significantly higher in M1 and lower in M2 macrophages, as opposed to control GFP_cyt_ that remained stable (Figure S5P). These findings indicated a reciprocal interaction whereby pre-existing GroEL2 variation influences macrophage polarization, and alterations in the host microenvironment, in turn, induce changes in GroEL2 at the single macrophage level.

**Figure 5.**
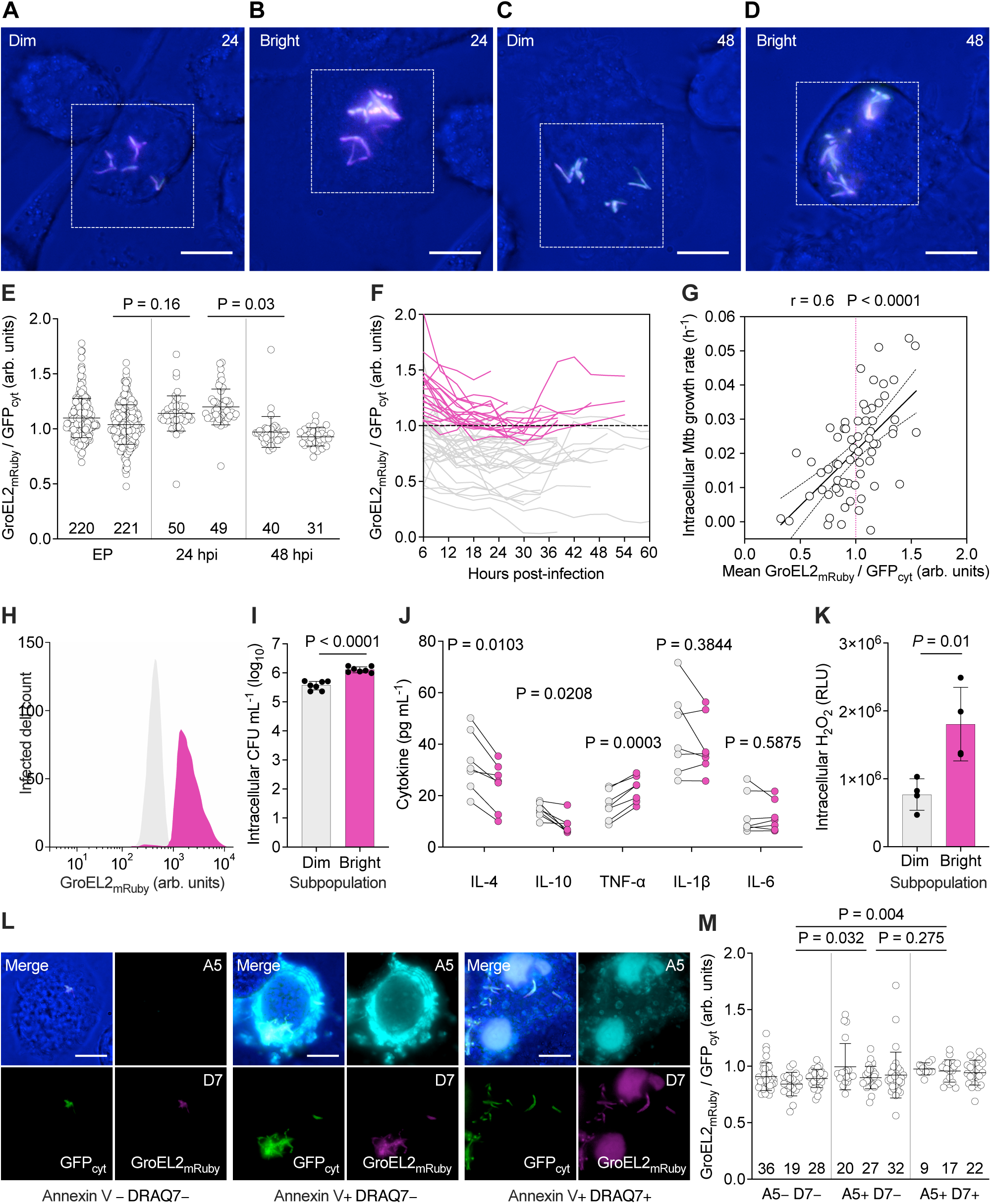
GroEL2_mRuby_ phenotypic variation predicts different macrophage responses **(A–D)** Snapshot images of THP-1 cells infected with GroEL2_mRuby_-GFP_cyt_ reporter at 24 and 48 hpi (n = 3). GroEL2_mRuby_-Dim (A, C) and -Bright (B, D) intracellular bacilli are shown (dashed boxes). Maximum intensity projections of four images from a z-stack of 8 µm; bright-field (blue), GFP_cyt_ (green), and GroEL2_mRuby_ (magenta) channels are merged. Scale bars, 10 µM. (E) GroEL2_mRuby_ normalized to GFP_cyt_ mean fluorescence of exponential-phase (EP) bacilli and intracellular infectious foci at 24 and 48 hpi. Lines represent mean ± SD (n = 2, with cell number shown at the bottom of the graphs). Significance by nested one-way ANOVA (P = 0.034) followed by Tukey multiple comparisons test (adjusted P values). (F) Time course of GroEL2_mRuby_/GFP_cyt_ ratio for single infectious foci, classified as Dim (mean ratio < 1, n = 31) or Bright (mean ratio > 1, n = 20). Dashed line indicates the threshold ratio. Data are from three independent experiments. (G) Pearson’s correlation between intracellular growth rate and GroEL2_mRuby_/GFP_cyt_ ratio of individual intracellular infectious foci over the infected cell lifetime (n = 64 from two independent experiments). The dotted line indicates the threshold ratio. Simple linear regression fit (solid line, R^2^ = 0.36, P < 0.0001) and 95% confidence bands (dashed lines). (H) Flow cytometry analysis of infected THP-1 macrophages at 24 hpi, sorted into GroEL2_mRuby_-Dim (gray) and -Bright (pink) subpopulations based on intracellular fluorescence. Representative overlays are normalized to 10,000 events (n = 7 independent experiments). (I) Bacterial load in GroEL2_mRuby_-Dim and -Bright THP-1 subpopulations at 24 hpi. Lines represent mean ± SD (n = 7). Significance by two-tailed unpaired *t*-test (95% confidence level). (J) Immunoluminometric quantification of cytokines at 48 hpi in GroEL2_mRuby_-Dim (gray) and -Bright (pink) THP-1 subpopulations sorted at 24 hpi as in (n = 7). Significance by multiple paired *t*-tests with Holm-Sidak correction (adjusted P values). (K) Quantification of oxidative burst in THP-1 in subpopulations sorted at 48 hpi based on intracellular GroEL2_mRuby_ levels. Lines represent mean ± SD (n = 4 independent experiments). Significance by two-tailed unpaired *t*-test (95% confidence level). (L) Representative snapshot images of THP-1 cells infected with GroEL2_mRuby_-GFP_cyt_ reporter at 48 hpi, stained with annexin V (A5) and DRAQ7 (D7) (n = 3 independent experiments). Maximum intensity projections of four images from a z-stack of 6 µm; bright-field (blue), A5 (cyan), GFP_cyt_ (green), GroEL2_mRuby_ and D7 (magenta) channels are merged. Scale bars, 10 µm. (M) GroEL2_mRuby_/GFP_cyt_ ratio for single intracellular infectious foci at 48 hpi in THP-1 macrophages, classified according to A5 and D7 negative (–) or positive (+) staining. Lines represent mean ± SD (n = 3, with cell number shown at the bottom of the graphs). Significance by two-way ANOVA (P = 0.009), followed by multiple comparisons with two-stage linear step-up procedure of Benjamini, Krieger and Yekutieli (individual P values).

Consistent with different macrophage activation states relative to GroEL2_mRuby_, we found that cells infected with GroEL2_mRuby_-Bright bacilli produced more reactive oxygen species than those infected with GroEL2_mRuby_-Dim bacilli at 48 hpi (Figure 5K). Thus, we asked whether these macrophage subpopulations underwent distinct forms of cell death (Ding and Briken, 2026). We stained infected macrophages with annexin V, an early marker of apoptosis, and with DRQ7, a marker of membrane permeabilization. At 48 hpi, 40% of macrophages were negative for both markers, 37% were positive for annexin V only, and 23% were dual-positive, suggesting necrosis secondary to apoptosis (Figure 5L). Importantly, bacilli infecting apoptotic or necrotic macrophages expressed significantly higher GroEL2 levels compared to those in viable macrophages (Figure 5M).

In summary, our findings demonstrate that pre-existing GroEL2 phenotypic variation influences early host polarization events and is dynamically regulated during infection. Higher GroEL2 expression increases the likelihood of triggering extrinsic apoptosis and secondary necrosis, processes associated with host-protective bacterial containment and host-detrimental bacterial spread, respectively. Early diversification of macrophage responses, contingent on *M. tuberculosis* phenotypic variation, may ultimately contribute to lesion heterogeneity (Flynn and Chan et al., 2022).

## GroEL2 triggers NF-κB and IRF signaling pathways with potential for anti-inflammatory therapy

Different PRRs have been implicated in GroEL2 recognition, including TLR2 and TLR4 (Daher et al., 2023). TLRs bind microbial ligands through their ectodomains, inducing TIR domain dimerization and recruitment of adaptor proteins into cytosolic supramolecular organizing centers, namely the myddosome and the triffosome (Fitzgerald and Kagan, 2020). These signaling hubs initiate transcriptional programs that regulate innate antimicrobial defenses, cytokine secretion, antigen presentation, and influence adaptive immunity. TLRs on the plasma membrane are mainly involved in MyD88-dependent myddosome assembly, whereas endosomal receptors, which can be recruited to phagosomes, can assemble either MyD88-dependent myddosomes or TRIF-dependent triffosomes in the case of TLR3 and TLR4 (Fitzgerald and Kagan, 2020). The myddosome drives NF-κB-mediated inflammatory responses and is essential for efficient host control of tuberculosis (Sánchez et al., 2010; Hayden and Ghosh, 2011). In contrast, the triffosome activates interferon regulatory factors (IRF), inducing type I interferons (IFN-I), which can impair host control of *M. tuberculosis* (McNab et al., 2015). While TLR2 heterodimers act exclusively at the plasma membrane, TLR4 dimers are unique in transmitting signals both from the plasma membrane and from endosomal compartments upon ligand-induced receptor endocytosis (Fitzgerald and Kagan, 2020). Hence, TLR4 induces early NF-κB activation from the plasma membrane, followed by IRF activation and a late wave of NF-κB activation from recycling endosomes and phagosomes. Improved understanding of these events is critical to unravel early host–pathogen interactions.

To investigate TLR signaling induced by GroEL2 in human macrophages, we stimulated wild-type (WT) and MyD88 knockout (KO_MyD88_) dual-reporter THP-1 cells, which simultaneously monitor NF-κB and IRF expression, with untagged recombinant GroEL2 verified for low endotoxin levels (Figures S6A and S6B). GroLE2 significantly activated NF-κB in WT cells after 24 hours (Figures S6C and S6D). While antibody inhibition of TLR2 had no effect against GroEL2 stimulation, chemical inhibition of TLR4 with CLI-095 significantly decreased NF-κB activation (Figure S6D). Additionally, low but significant NF-κB activation was also detectable in KO_MyD88_ cells and inhibited by CLI-095, consistent with a MyD88-independent mechanism contingent on TLR4. Consistent with this, GroEL2 activated IRF in both WT and KO_MyD88_ cells, and CLI-095 significantly inhibited these responses, indicating triffosome-dependent signaling from endosomal compartments (Figure S6E). Thus, our data support the involvement of TLR4 in GroEL2-induced signaling in human macrophages, as previously reported in murine cells (Bulut et al., 2005; Cehovin et al., 2010).

Next, we asked whether this mechanism occurred similarly during infection. We transcriptionally silenced *rv2224c* in *M. tuberculosis* (*rv2224c*_si_), leading to strong accumulation of the uncleaved GroEL2_H_ form into the supernatant, as opposed to control bacteria (CT), which released lower amounts of both GroEL2 variants (Figure 6A). As early as 6 hpi, GroEL2_H_ caused low but significant activation of both NF-κB and IRF reporters, with the latter being similarly activated also in KO_MyD88_, consistent with endosomal signaling (Figures S6F and S6G). At 24 hpi, NF-κB and IRF reporters were significantly activated in both WT and KO_MyD88_ macrophages, confirming the implication of MyD88-dependent and -independent pathways in response to GroEL2_H_ (Figures 6B and 6C). While NF-κB activation was significantly stronger in WT cells, IRF activation was stronger in KO_MyD88_ cells, possibly suggesting increased TLR4 endocytosis and delivery of TLR4 pools to phagosomes in cells devoid of MyD88. Indeed, we detected TLR4 inhibition by CLI-095 only in KO_MyD88_ cells, with a significant decrease in both NF-κB and IRF activation. These results support the involvement of TLR4 in GroEL2_H_-mediated IRF stimulation and suggest crosstalk between cellular compartments; however, they do not exclude contributions from additional host receptors.

**Figure 6.**
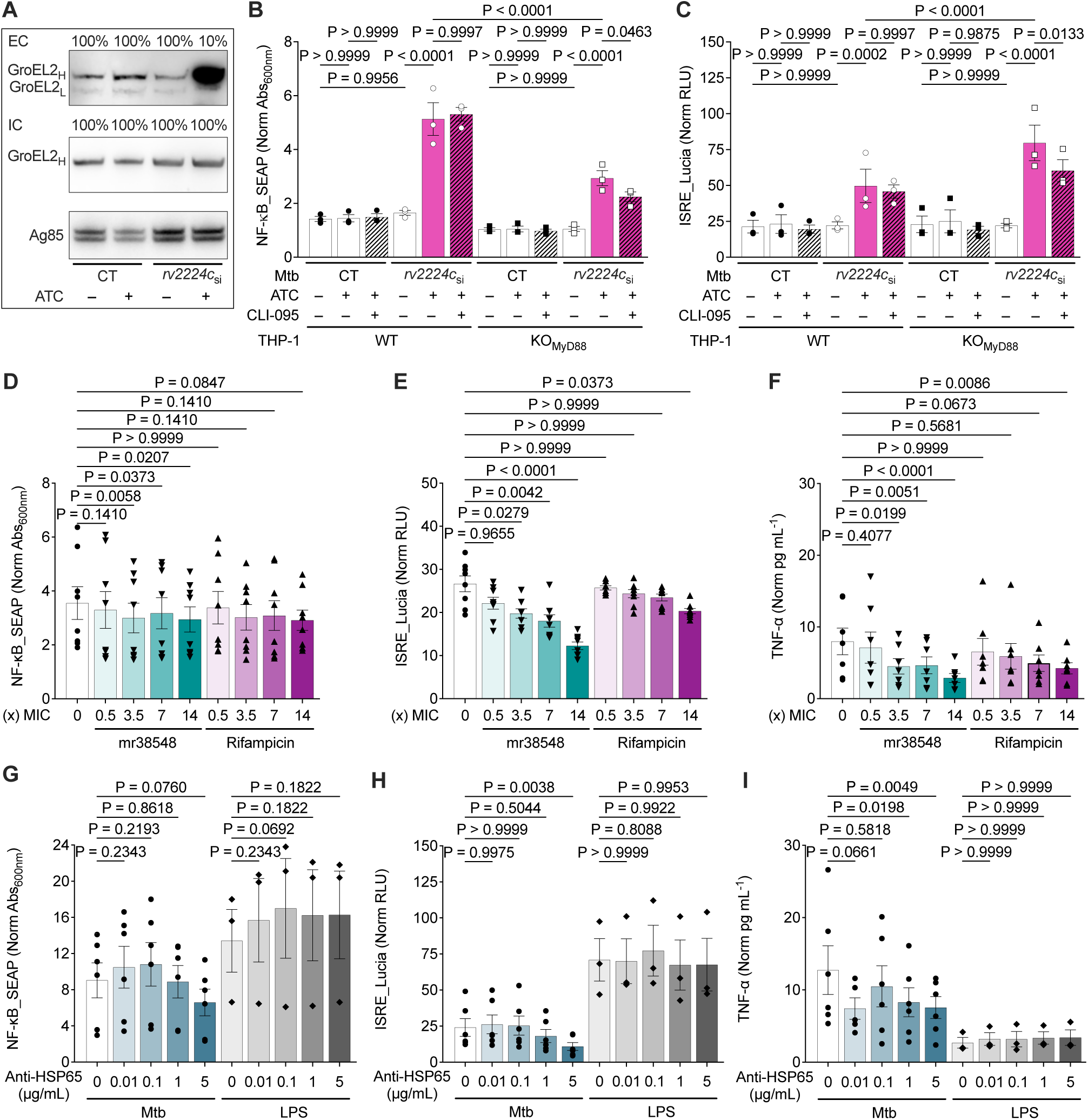
GroEL2_H_ triggers NF-κB and IRF signaling and can be targeted to decrease inflammation **(A)** Immunoblotting of *M. tuberculosis* extracellular (EC) and intracellular (IC) fractions using anti-HSP65 mouse monoclonal IgG_2a_. Control (CT) and *rv2224c*-silenced (si) strains in the absence (–) or presence (+) of ATC inducer (200 ng/mL). Loaded sample amounts in percentage, high (H) and low (L) molecular-weight GroEL2 bands, and Ag85 as a control are indicated. **(B and C)** Activation of NF-κB (B) and IRF (C) in WT (circles) and KO_MyD88_ (squares) dual-reporter THP-1 cells 24 hpi. Cells were infected with either *M. tuberculosis* CT (filled symbols) or *rv2224c*_si_ (open symbols) at MOI 2 for 2 hours, 4 days after induction of silencing. Absence (–) or presence (+) of inducer and TLR4 inhibitor (760 nM) is indicated. NF-κB_SEAP (B) and ISRE_Lucia (C) signals are normalized to uninfected cells. Significance by one-way ANOVA (P < 0.0001), followed by Šidák’s multiple comparisons test (adjusted P values). Lines represent mean ± SEM (n = 3). **(D–F)** Activation of NF-κB (D) and IRF (E), and TNF-α production (F) in WT THP-1 dual-reporter cells infected with *M. tuberculosis* H37Rv (MOI 5 or 10) for 3 hours. Cells were left untreated (circles) or treated with mr38548 (MIC: 0.88 µg/mL; inverted triangles) or RIF (MIC: 0.076 µg/mL; triangles) at different concentrations relative to the MIC (x) for 24 hours. Values are normalized to uninfected and untreated cells. Significance by non-parametric Friedman test: P = 0.016 (D) and P < 0.0001 (E and F), followed by Dunn’s multiple comparisons test (adjusted P values). Lines represent mean ± SEM (n ≥ 6). **(G–I)** Activation of NF-κB (G) and IRF (H), and TNF-α production (I) in WT THP1 dual-reporter cells infected with *M. tuberculosis* H37Rv (MOI 5 or 10; circles) or stimulated with LPS (100 ng/mL; diamonds) for 5 hours. Cells were left untreated or exposed to different concentrations of monoclonal anti-HSP65 IgG_2a_. Values are normalized to uninfected and untreated cells. Significance by mixed-model one-way ANOVA: P = 0.0014 (G), P = 0.0002 (H), and P = 0.0274 (I), followed by Šidák’s multiple comparisons test (adjusted P values). Lines represent mean ± SEM (n ≥ 3).

To start searching for GroEL2 targets in macrophages, we performed preliminary immunoprecipitation coupled with mass spectrometry in WT_MyD88_ and KO_MyD88_ THP-1 cells using His-tagged recombinant GroEL2. Differential analysis revealed host proteins potentially associated with GroEL2 (Table S4). In WT cells, GroEL2 pull-down was enriched in proteins involved in NF-κB signaling, actin dynamics, cytoskeletal remodeling, phagosome maturation, and apoptosis (Figure S6H). In contrast, GroEL2 pull-down in KO_MyD88_ cells was mainly enriched with proteins involved in lipid metabolism. Although these results do not demonstrate direct protein-protein interactions, they point out candidate host pathways potentially affected by GroEL2.

Next, we asked whether targeting GroEL2 could impact innate immune signaling. We synthesized mr38548, a small molecule previously reported as a putative GroEL2 inhibitor (Sarkar et al., 2022). Despite the weak affinity of mr38548 for recombinant GroEL2 in vitro (Figure S7A), we confirmed its intracellular activity, which was about tenfold lower than that of RIF (Figure S7B). We infected macrophages with WT *M. tuberculosis* at high multiplicity of infection (MOI) and treated them either with mr38548 or with RIF. Consistent with our hypothesis, mr38548, but not RIF, caused a significant, dose-dependent decrease in NF-κB and IRF activation, as well as in TNF-α secretion (Figures 6D–6F), indicating that GroEL2 targeting rather than mere bacterial killing was causing a reduction in inflammation (Figure S7B). We further confirmed the specificity of mr38548 against *M. tuberculosis*, as LPS-induced inflammation was not altered by either mr38548 or RIF treatment (Figures S7C–S7E). Consistent with these results, targeting GroEL2 with a monoclonal antibody significantly reduced IRF activation and TNF-α production during *M. tuberculosis* infection, but not in response to LPS, albeit to a lesser extent than chemical inhibition (Figures 6G–6I).

In conclusion, our findings indicate that cell-to-cell variation in GroEL2 expression influences both bacterial fitness and early macrophage responses. Bacilli with low GroEL2 levels primarily cause moderate IRF activation, leading to increased production of anti-inflammatory cytokines and likely IFN-I. The latter can signal through the interferon-α/β receptor (IFNAR), promoting pathogen-permissive M2 macrophage polarization. In contrast, bacilli with high GroEL2 levels, particularly exporting the multimeric variant, cause strong activation of both NF-κB and IRF and of late NF-κB signaling. Consequently, NF-κB increases TNF-α production, which can further amplify NF-κB via the TNF receptor (TNFR), promoting pathogen-restrictive M1 macrophage polarization. Although additional mechanistic details remain to be clarified, such as the specific roles of GroEL2 oligomeric variants, contributions from other PRRs, and spatiotemporal dynamics of cell signaling, endosomal trafficking, and phagosomal maturation, our results position GroEL2 as a key modulator of bacterial adaptation and host immunity. Therefore, targeting GroEL2 has strong therapeutic potential to decrease excessive inflammatory responses without compromising drug-mediated intracellular control of *M. tuberculosis* and may help clear the infection faster (Figure 7).

**Figure 7.**
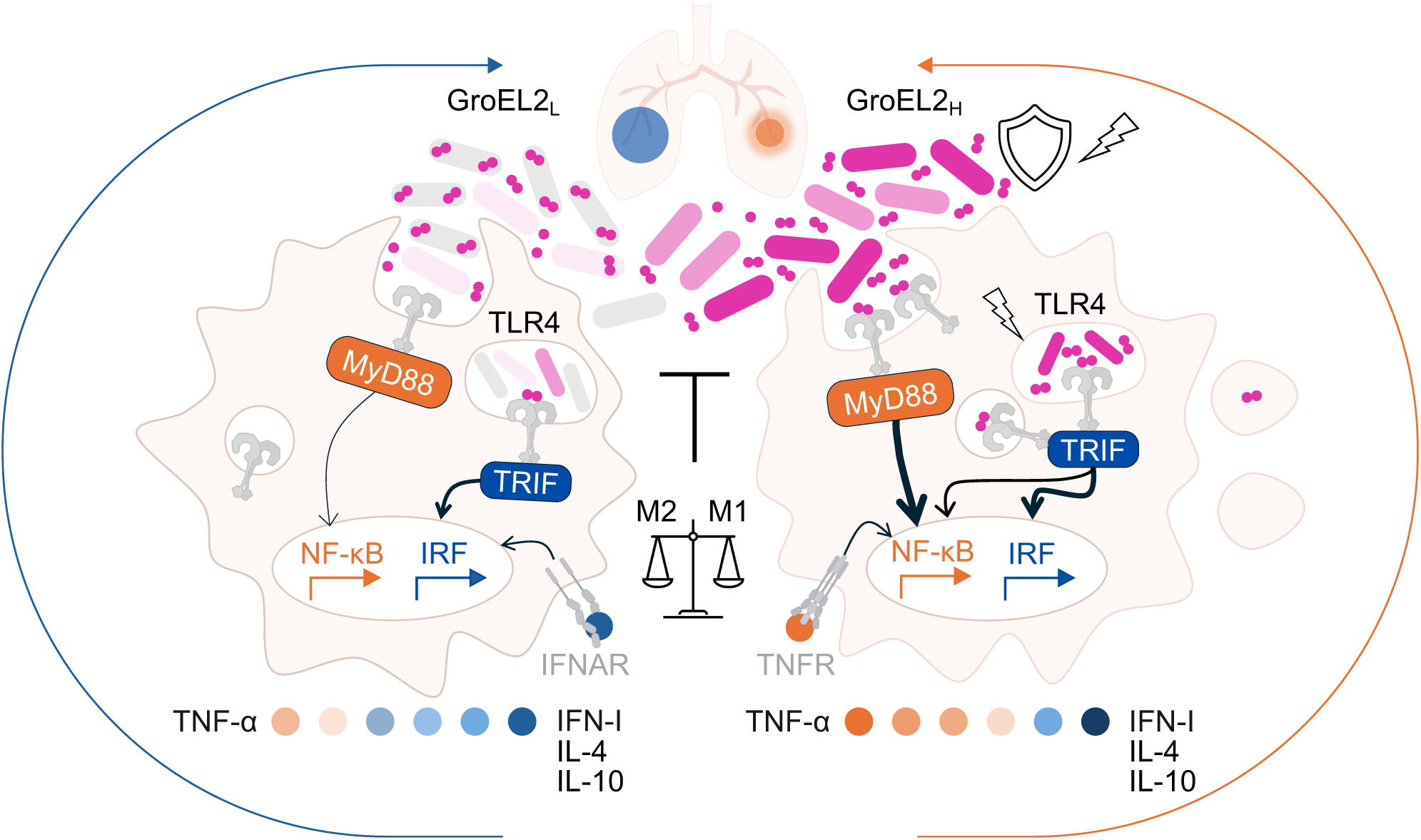
Schematic illustration of the effect of GroEL2 heterogeneity on single-cell infection. Clonal *M. tuberculosis* cells heterogeneously expressing GroEL2 (gray to pink gradient) interact with individual macrophages, leading to distinct possible outcomes. GroEL2-Dim bacilli (gray and light pink) express lower levels of GroEL2 (left), which mainly localizes as intracellular foci and is poorly released into the extracellular milieu as equal amounts of different oligomeric forms. This induces weak NF-κB and relatively higher IRF activation, resulting in increased production of anti-inflammatory cytokines and type I IFNs (blue circles) compared with pro-inflammatory TNF-α (orange circles). This response favors a growth-permissive M2 macrophage phenotype and may contribute to setting the initial conditions for future lung lesion expansion (blue arrow and circle). GroEL2-Bright bacilli (dark pink) express and release higher levels of multimeric GroEL2 (right), which strongly activates NF-κB and IRF, amplifying TNF-α production and favoring a growth-restrictive and pro-inflammatory M1 macrophage phenotype. This response may contribute to setting the initial conditions for future lung lesion containment (orange arrow and circle). The thickness of black arrows reflects the intensity of signaling, which involves both the plasma membrane and endosomal or phagosomal compartments, with TLR4 involvement. Other host receptors likely contribute to amplifying single-cell responses, such as IFNAR and TNFR. GroEL2 inhibition may support pathogen control, decrease pro-inflammatory deregulation, and limit lung pathology.

## Discussion

Central obstacles to the control of tuberculosis are the poor predictability of *M. tuberculosis* infection outcomes, disease progression, and response to treatment. Clinical manifestations can vary markedly among and within individuals. This variability reflects the pathogen’s ability to fine-tune its physiology to complete a successful life cycle across infection, immune evasion, persistence, pathology, and transmission (Ehrt et al., 2018; Simmons et al., 2018; Chandra et al., 2022; Behr et al., 2024; Dheda et al., 2025). The changeable physiology of *M. tuberculosis* is rooted in its inherent and stress-triggered phenotypic variation in key processes, such as DNA replication, growth rate, energy production, carbon and lipid metabolism, redox homeostasis, and antigen secretion (Dhar et al., 2016; Chung et al., 2022; Sherry and Rego, 2024). All these processes are essential for adaptation (Wilburn et al., 2018; Pacl et al., 2018; Mekonnen et al., 2021; Mishra et al., 2021; Nargan et al., 2024) and ultimately contribute to clinical heterogeneity (Flynn and Chan, 2022; Lavin and Tan, 2022; Scriba et al., 2024). Understanding how this phenotypic diversification arises and translates into cell-to-cell functional differences in increasingly complex host microenvironments can help identify biomarkers and targetable pathways to improve and personalize interventions.

In vivo, *M. tuberculosis* is confronted with hostile conditions, such as fluctuations in carbon sources, pH, oxygen tension, reactive species, and antibiotics, which cause potentially lethal stress that requires effective countermeasures (Pacl et al., 2018). To this end, *M. tuberculosis* relies on several sensing and detoxification functions, including redox-sensitive Fe-S cluster proteins, which monitor microenvironmental redox changes and maintain the electron transport chain, supporting metabolic and physiological adaptations required for survival (Mishra et al., 2021; Berdal et al., 2026). In this study, we use FdxA, a ferredoxin upregulated under different redox stressors in vitro and in vivo (Schnappinger et al., 2003; Kendall et al., 2004; Kesavan et al., 2009), as a starting marker of stress responsiveness in *M. tuberculosis*. We show that phenotypic variation in FdxA predicts subpopulation-specific RNA signatures related to differences in regulatory, metabolic, respiratory, and adaptive pathways. Not surprisingly, several heat-shock proteins, mainly DnaK, ClpB, and GroEL2, are upregulated in highly stress-responsive *M. tuberculosis* subpopulations.

Molecular chaperones are oligomeric machineries with a cavity that aids de novo folding and stress-induced refolding of proteins (Balchin et al., 2020). GroEL2 is essential for *M. tuberculosis* optimal growth, acts independently of the GroEL1 co-chaperonin GroES, and has poor ATPase activity but has thioesterase activity on short acyl-CoA substrates, possibly contributing to metabolic plasticity and cell detoxification (Qamra et al., 2005; Fan et al., 2012; Chilukoti et al., 2016; Zhou et al., 2026). It was reported that the *Escherichia coli* ortholog GroEL, which, however, cannot be functionally complemented by *M. tuberculosis* GroEL2 (Chilukoti et al., 2016), exhibits diffused cytosolic distribution as well as colocalization with the cell division protein FtsZ (Charbon et al., 2011). Biochemical reconstitution and electron microscopy analysis revealed that GroEL recognizes ribosome-bound nascent proteins and favors de novo co-translational folding with GroES (Roeselová et al., 2025). These findings suggest spatially distinct roles for GroEL in *E. coli*. In *M. tuberculosis*, GroEL2 is thought to be membrane associated, and it was identified in whole cell extract, membrane fraction, and culture filtrate (de Souza et al., 2011; Birhanu et al., 2019) as well as in mycobacterial extracellular vesicles (Palacios et al., 2021). This implies that GroEL2 can carry out different functions inside and outside the cell, albeit the export mechanism remains unknown.

Here, using fluorescently labeled GroEL2, we observe considerable cell-to-cell heterogeneity in *M. tuberculosis*, even under optimal growth conditions. We show not only uneven expression levels but also distinct patterns of localization. While exponentially growing *M. tuberculosis* cells mainly exhibit transient localization foci typically at the cell poles, upon drug, oxidative, and nutritional stress, GroEL2 can evenly delocalize throughout the cell, relocalize to the nucleoid, form patchy foci, or move along the cell boundary. Distinct cellular locations can suggest diverse functions associated with cytosol, DNA, RNA, ribosomes, or cell envelope dynamics (Dhar et al., 2016; Chung et al., 2022). In *M. tuberculosis*, the chaperone ClpB was previously shown to form fluorescent patches that colocalize with damaged proteins upon drug exposure, suggestive of a reparative function (Vaubourgeix et al., 2015). In *E. coli*, chaperone-dependent removal of aggregated proteins implicated in energy production was proposed to induce exit from dormancy (Bollen et al., 2025). Likewise, an anti-aggregative role was reported for *M. tuberculosis* GroEL1 and GroEL2 in vitro (Qamra et al., 2005; Shahar et al., 2011; Fan et al., 2012; Chilukoti et al., 2016). However, probably due to its unstable oligomeric nature, GroEL2 prevents protein aggregation more efficiently at higher concentrations. Consistent with this, we show that cells inherently expressing higher GroEL2 levels grow faster and are more tolerant to stressful conditions. Additionally, GroEL2 is induced in response to different antimicrobial stressors, leading to a rise in the extracellular GroEL2_H_ multimeric variant. Context-specific spatiotemporal patterns of GroEL2 expression in *M. tuberculosis* are likely associated with its tight regulation at transcriptional (Stewart et al., 2002; Stapleton et al., 2012), post-transcriptional (Houghton et al., 2021), and post-translational levels (Naffin-Olivos et al., 2014; Birhanu et al., 2019; Zhou et al., 2026). This multilayer control also reflects a fundamental paradox in *M. tuberculosis* and, more broadly, in pathogens’ biology. On the one hand, molecular chaperones support pathogen survival by promoting protein homeostasis and stress adaptation. On the other hand, they facilitate binding, invasion, and activation of antigen-presenting cells, which elicit innate responses and T cell priming, thus contributing to pathogen control (Henderson et al., 2006). Consistent with this, deletion of the transcriptional repressor HspR, which leads to upregulation of several chaperones, including DnaK, GrpE, DnaJ, and ClpB, was shown to impair *M. tuberculosis* survival in a chronic murine infection model, confirming their role in immune activation and the need to be fine-tuned to enable pathogen persistence (Stewart and Young, 2004). Similarly, lack of proteolytic cleavage by Hip1, resulting in increased GroEL2_H_ multimeric form, was shown to induce pro-inflammatory cytokines in murine macrophages, possibly based on TLR2 and MyD88-dependent signaling, and to support disease control in mice (Sia and Rengarajan, 2019; Madan-Lala et al., 2011; Naffin-Olivos et al., 2014). Additionally, dendritic cells stimulated with the multimeric variant exhibited greater antigen presentation to naïve CD4+ T cells than the cleaved variant (Georgieva et al., 2018). Importantly, mice lacking TLR2 or MyD88 and humans with polymorphisms in the corresponding genes have increased susceptibility to tuberculosis. Likewise, TLR4-defective C3H/HeJ mice control *M. tuberculosis* pulmonary burden less efficiently and do not respond to human HSP60, and TLR4 polymorphisms are associated with greater tuberculosis severity in certain human populations (Mortaz et al., 2015). However, while TLR2 primarily senses mycobacterial lipoproteins, TLR4 mainly recognizes mycobacterial and human chaperones or derived peptides (Ohashi et al., 2000; Bulut et al., 2005; Jo et al., 2007; Cehovin et al., 2010; Vila-Casahonda et al., 2025).

In this study, we show that transcriptional silencing of *rv2224c* (Hip1) causes significant extracellular release of multimeric GroEL2_H_, which activates not only NF-κB, as expected, but also IRF pathways, the latter even more robustly in KO_MyD88_ macrophages. These findings provide the first link between GroEL2 and MyD88-independent signal transduction, coordinated between the plasma membrane and intracellular compartments, supporting TLR4 implication (Fitzgerald and Kagan, 2020; Wang et al., 2024). However, while chemical inhibition of TLR4 decreased NF-κB and IRF activation to background levels after stimulation with recombinant GroEL2, the effect was less pronounced during infection and apparent only in KO_MyD88_ macrophages. This has at least two non-mutually exclusive explanations. First, excessive levels of GroEL2_H_ can strongly induce TNF-α, thus amplifying NF-κB activation via TNFR (Hayden and Ghosh, 2011; Chen et al., 2023). Second, other plasma membrane receptors, such as LOX-1 (Vinod et al., 2021) and most likely CD43, which binds GroEL2 and facilitates *M. tuberculosis* uptake and TNF-α production (Torres-Huerta et al., 2017), can further contribute to NF-κB background signaling, masking TLR4 inhibition. Additionally, it was reported that the TLR4/TLR2 ratio is higher in M1 macrophages, whereas TLR4 deficiency promotes M2 polarization in adipose tissue macrophages (Wang et al., 2014). At present, we cannot exclude that macrophage variation in TLRs expression also contributes to the observed patterns of cell signaling activation induced by GroEL2. These results raise additional questions about how this antigen is recognized, transported, and acts at the intersection of innate and adaptive immunity.

Preliminary co-immunoprecipitation coupled with mass spectrometry has already enabled us to identify candidate host proteins that may interact with GroEL2. The most significantly enriched factors included FAM193B, previously associated with MAPK/ERK and PI3K/AKT pathways (Xie et al., 2021); paxillin and the Rho GTPase ARHGEF40, implicated in cytoskeletal remodeling and phagosome dynamics (Bhattacharya et al., 2025; Wang et al., 2017); and Ca²⁺/calmodulin-dependent kinases involved in phagosome maturation (Vergne et al., 2004). We also detected TECR, involved in the synthesis of sphingolipids that influence membrane integrity, actin dynamics, *M. tuberculosis* phagocytosis, and regulation of inflammation and apoptosis (Niekamp et al., 2021); TECPR1, which recognizes cytosolic sphingomyelin and recruits LC3 to damaged endomembranes (Boyle et al., 2023); and pro-apoptotic proteins, such as the transcription factor BCLAF1 (Shao et al., 2016). Overall, these results imply that GroEL2 participates in intracellular trafficking events and influences the metabolism of bioactive lipids and host cell death pathways. However, whether these effects take place via GroEL2 active or passive mechanisms remains to be determined. In future work, we will probe macrophage transcriptomic, proteomic, and in-cellulo responses to *M. tuberculosis* expressing either GroEL2_H_ or GroEL2_L_, aiming to further clarify their role in host-cell modulation and endosomal trafficking (Zuppini et al., 2025).

Apoptosis promotes phagocytic clearance, limits bacterial replication, dampens inflammation, and favors antigen presentation, whereas necrosis promotes bacterial dissemination, exacerbates inflammation, and leads to immunopathology (Ding and Briken, 2026). GroEL2 was previously reported to inhibit apoptosis via the mitochondrial protein mortalin (Joseph et al., 2017); conversely, we find that macrophages with higher intracellular GroEL2 levels are more likely to undergo apoptosis, and those with the highest levels undergo necrosis, suggesting a continuum from host-protective to host-detrimental cell death pathways. This is consistent with the positive correlation we observe between intracellular GroEL2 levels, bacterial burden, and pro-inflammatory responses in macrophages. Thus, we posit that pre-existing GroEL2 heterogeneity is associated with a spectrum of innate responses, which in turn influence GroEL2 intracellular dynamics, collectively contributing to early infection outcomes, with likely consequences for lesion formation and evolution (Figure 7). We propose that fine-tuning GroEL2 may convert tissue-damaging pathology into host-protective responses, and vice versa, with clinical implications for disease heterogeneity, providing a rationale for more targeted interventions (Chen et al., 2023; Scriba et al., 2024; Sweet et al., 2025).

Our pilot study in patients’ sputum shows the clinical potential of GroEL2 as a positive biomarker of persistent bacilli in subclinical cases and during treatment. Our findings are consistent with GroEL2 downregulation in sputum from active tuberculosis patients (Honeyborne et al., 2016) and with upregulation in sputum from patients for whom therapy is ineffective (Sharma et al., 2017; Shaikh et al., 2021). Overall, this supports the hypothesis that GroEL2 promotes *M. tuberculosis* survival in vivo under restrictive immune and drug pressure. However, to validate GroEL2 as a prognostic and predictive biomarker, a larger prospective clinical study will be required, including geographically diverse cohorts and possibly other specimen types, such as bronchoalveolar lavages or biopsies, which can provide higher-quality samples and insights into antigen spatial distribution and heterogeneity (Nargan et al., 2024). Additionally, given the high conservation of GroEL2 among mycobacterial species (Yang et al., 2012), molecular, immunological, or spectroscopy-based approaches (Meehan et al., 2019; Stöckel et al., 2017) should be compared to enable species-specific early diagnosis and to inform patient stratification and personalized therapy (Pai et al., 2023). In the need for original and fast-acting anti-tubercular strategies (Dartois et al., 2025), various immunotherapeutic approaches have gained momentum (Chen et al., 2023; Sweet et al., 2025; Vila-Casahonda et al., 2025). However, targeting or modulating host immune pathways requires careful assessment of potential secondary effects. For instance, inhibition of pathways critical for anti-mycobacterial immunity, as with TNF-α antagonists or JAK inhibitors, decreases inflammation but can favor tuberculosis progression or relapse (Chen et al., 2023; Brehm et al., 2024; Sweet et al., 2025). Achieving the right inflammatory balance could enhance pathogen clearance while preventing tissue pathology (Figure 7). Targeting chaperones is a viable approach, also thanks to the relatively low conservation between mycobacterial and human orthologs (Kampinga et al., 2009). For instance, inhibition of *M. tuberculosis* DnaK (HSP70) was reported to potentiate antibiotics and prevent drug resistance (Hosfelt et al., 2022). In the present study, we show that targeting GroEL2 not only impairs *M. tuberculosis* but also moderately decreases inflammation, possibly helping to control bacterial burden while limiting tissue pathology and transmission. Successful examples of targeting pathogens with positive immunomodulatory effects have been reported for pathogenic *E. coli*, *Pseudomonas aeruginosa*, and *M. tuberculosis* (Parveen et al., 2023; Simonis et al., 2023; Wang et al., 2023). Furthermore, we showed that decreasing phenotypic variation in a stress-response process, which we refer to as pheno-tuning, increases *M. tuberculosis* susceptibility to standard therapy (Mistretta et al., 2024). Our future efforts will include in silico exploration of more specific GroEL2 covalent inhibitors and biotherapeutics, as well as optimization of GroEL2 pheno-tuning, aiming both to undermine *M. tuberculosis* fitness and to boost the innate antimicrobial capacity of the host cell to clear the infection, reducing the risk for tissue damage.

In conclusion, this work demonstrates that focusing on *M. tuberculosis* phenotypic variation at the host–pathogen interface represents a promising source of diagnostic biomarkers and therapeutic targets, which may aid in designing future precision medicine strategies.

## Acknowledgments

We thank Pierre-Henri Commere (Flow Cytometry Platform, Institut Pasteur, Paris, France) for his advice on flow cytometry and cell sorting. We thank Audrey Hessel (Production and Purification of Recombinant Proteins Technological Platform, Institut Pasteur, Paris, France) for her valuable assistance with GroEL2 protein purification. We thank Yanis Dahoumane, Lénaig Le Fouler, and Nathalie Jolly (Clinical Research Coordination Office, Institut Pasteur, Paris, France) and Minerva Cervantes Gonzalez (Infectious Diseases Department, Bichat University Hospital, AP-HP, Paris, France) for their support in establishing the clinical study collaboration. We thank Tri Tho Nguyen for advice on the preparation of the hydrogel used for time-lapse imaging of infected macrophages.

RNA sequencing was performed by the ICGex NGS platform of the Institut Curie supported by the French National Research Agency grants ANR-10-EQPX-03 and ANR-10-INBS-09-08, by the Canceropole Ile-de-France, and by the SiRIC Grant INCa-DGOS- 4654. The Image Analysis Hub is part of the France-BioImaging research infrastructure (https://ror.org/01y7vt929), supported by the French National Research Agency (ANR-24-INBS-0005 FBI BIOGEN).

This work was mainly supported by the following grants: French National Research Agency grants ANR-10-LABX-62-IBEID and ANR-17-CE11-0007-01 PersisTB to G.M.; Institut Carnot Pasteur Microbes & Health Emergence Project INNOV-68-20 to G.M. and C.C.; French National Research Agency grant ANR-21-CE15-0045 TREATABLE to G.M. and C.R.; Institut Pasteur EID Junior call 2023 to L.P., IMI 2 Joint Undertaking Grant 853989, receiving support from the European Union’s Horizon 2020 research and innovation programme and EFPIA and Global Alliance for TB Drug Development non-profit organization, Bill & Melinda Gates Foundation, University of Dundee to G.M., and Institut Pasteur core funding to G.M.

## Author contributions

Conceptualization: G.M.; Methodology: L.P., L.K.S., J.C.Z.A., M.P., O.H., C.L., A.L., E.B., and G.M.;

Software: M.A., and J.Y.T.; Validation: L.P., L.K.S., N.G., M.P., N.P., E.P., M.C., E.K., C.R., and G.M.;

Formal Analysis: L.P., G.M., C.C., N.P., E.P., L.K.S., M.P., M.M., J.C.Z.A., and O.H. Investigation:

L.P., L.K.S., N.G., M.P., G.M., O.H., S.L., and Z.J.; Interpretation: L.P., G.M.; L.K.S., C.C., O.H., C.R.,

C.L., and A.L.; Resources: L.P., L.K.S., N.G., M.P., J.C.Z.A., O.H., S.L., and Z.J.; Writing − Original Draft: G.M.; Writing – Review & Editing: all authors; Visualization: L.P, G.M., C.C., E.K., and O.H.; Supervision: G.M., C.R., S.L.G., and N.G.; Project Administration: G.M.; Funding Acquisition: G.M., C.R., L.P, and C.C.

## Resources availability

### Lead contact

Further information and requests for resources and reagents should be directed to and will be fulfilled by the lead contact, Giulia Manina.

## Materials availability

Plasmids and bacterial strains generated in this study and the compound mr38548 are available from the lead contact with a completed Materials Transfer Agreement. Resources used in this study are listed in Table S5.

## Data and code availability

We have no restriction on data availability. The datasets generated in the present study are either available in the article or publicly available in different repositories.

The subpopulation RNA-seq data have been deposited in NCBI’s Gene Expression Omnibus (Clough et al., 2024) and are accessible through GEO Series accession number GSE324777 (https://www.ncbi.nlm.nih.gov/geo/query/acc.cgi?acc=GSE324777).

The mass spectrometry proteomics data have been deposited to the ProteomeXchange Consortium via the PRIDE (Perez-Riverol et al., 2025) partner repository with the dataset identifier PXD075516. The custom Napari-Omnipose model and Python notebook used to analyze time-lapse microscopy videos are available on GitLab (https://gitlab.pasteur.fr/iah-public/napari-omnibacteria).

## Materials and methods

### Bacterial strains and growth conditions

DNA cloning and sequencing procedures were carried out in chemically competent *E. coli* TOP10 or *E. coli* BL21 (DE3) when cloning was followed by protein purification. *E. coli* transformants were grown in Luria–Bertani (LB) medium with appropriate selection: 100 µg/mL ampicillin, 50 µg/mL kanamycin, or 100 µg/mL hygromycin. Induction of His-tagged GroEL2 in *E. coli* BL21 (DE3) was achieved in LB medium with 1 mM IPTG at 19 °C overnight. *M. tuberculosis* Erdman or H37Rv WT and derived strains were cultured in Middlebrook 7H9 broth (BD) supplemented with 0.5% BSA, 0.2% glucose, 0.085% NaCl, 0.5% glycerol, and 0.05% Tween-80 (unless otherwise specified). Mycobacterial transformants were selected on Middlebrook 7H10 agar plates (BD) supplemented 10% OADC (BD) and 0.5% glycerol with appropriate selection: 20 µg/mL kanamycin, 50 µg/mL hygromycin, or 15 µg/mL kanamycin with 50 µg/mL hygromycin. Bacterial stocks were prepared from late exponential-phase cultures (OD_600_ = 1.0) obtained from single colonies; aliquots were supplemented with 15% glycerol and stored at −80 °C. Primary cultures were started from a glycerol stock diluted 1:100 and used only once. *M. tuberculosis* strains were grown in Middlebrook 7H9 broth with appropriate selection at 37 °C under shaking conditions (50 rpm) to mid-log phase (OD600 = 0.4–0.8). Primary cultures were used to inoculate secondary cultures for final experiments at mid-log phase, unless otherwise specified, and without antibiotic selection prior to bacterial sorting. Hartmans–de Bont minimal medium was used for nutrients starvation. Transcriptional silencing of *rv2224c* was achieved in M*. tuberculosis* secondary cultures by adding 100 ng/mL anhydrotetracycline (ATC) for four days prior to the final experiment, during which ATC was maintained.

## Cell lines and culture conditions

THP-1 human monocytes (ATCC TIB-202) were cultured in RPMI 1640 (Gibco) supplemented with 10% heat-inactivated fetal bovine serum (FBS) and 100 µg/mL penicillin–streptomycin (Pen-Strep). THP-1 Dual™ and THP-1 Dual™ MyD88 knock-out cells (InvivoGen) were grown in RPMI 1640 (Gibco) supplemented with 10% FBS as above, 25 mM HEPES, 100 µg/mL Pen-Strep, 100 µg/mL Normocin, 10 µg/mL Blasticidin, and 100 µg/mL Zeocin.

For all three cell lines, initial culture was performed in 10 mL growth medium containing 20% heat-inactivated FBS. Flasks were incubated upright at 37 °C in a humidified 5% CO_2_ atmosphere for 3 days before being laid flat and incubated for a further 3 days. At this point, 10 mL of standard medium with 10% FBS was added and cells were grown to a density of 1x10^6^ cells/mL before being split 1:20 for fewer than 20 passages. Stocks were prepared in fresh freezing medium containing 95% FBS and 5% DMSO at 2–5x10^6^ cells/mL after confirming the absence of mycoplasma contamination.

For each experiment, a stock was thawed at 37 °C, washed once in 10 mL medium, and inoculated into 30 mL of prewarmed complete RPMI in a T-175 flask. Cells were propagated at 37 °C in a humidified 5% CO_2_ atmosphere until reaching a density no greater than 1x10^6^ cells/mL, for up to four days. Monocytes were differentiated to an M0 macrophage-like state by incubation with 30 ng/mL phorbol 12-myristate 13-acetate (PMA, Sigma) for 48 hours, followed by removal of PMA and a 24-hour rest period in growth medium prior to infection. M0 macrophages were polarized to M1 or M2 states by incubation for 48 hours with 20 ng/mL human IFN-γ (Bio-Techne) and 10 pg/mL LPS (Sigma) for M1, or 20 ng/mL human IL-13 and IL-4 (Bio-Techne) for M2 activation. After the 48-hour incubation, cells were immediately infected.

## Human pilot study and ethical approval

Human sputum samples were obtained from the pilot study DetectTB (Institut Pasteur n° 2021-005), designed to identify mycobacterial biomarkers predictive of bacterial persistence. The study protocol, patient information sheet, and procedures were approved (n° 2022-02/IRB/2) on May 23, 2022, by the Institut Pasteur Institutional Review Board (IRB00006966, Institut Pasteur IRB #1). All procedures complied with French regulations and General Data Protection Regulation (GDPR). The collaboration between Institut Pasteur and Assistance Publique–Hôpitaux de Paris (AP-HP) was formalized through a Research Collaboration Contract (AP-HP reference n° VAL / 2022-705/01; Institut Pasteur reference n° 2021-005).

Residual sputum samples originally collected for routine care at Hôpital Bichat–Claude Bernard (Paris, France) were accessed for research only after patients were informed and confirmed no opposition (or objection). Associated clinical and microbiological data, including age, gender, bacterial load, drug resistance, and selected laboratory parameters, were pseudonymized, transferred via REDCap secure web application, and accessible only to authorized personnel, ensuring confidentiality.

The study targeted 39 adult participants (18–65 years): 9 tuberculosis-negative controls, 12 clinically active tuberculosis patients (9 sampled before and during or after intensive treatment), and 9 subclinical tuberculosis cases (IGRA-positive with no clinical symptoms). Inclusion criteria were the ability to produce about 10 mL of sputum, weight 35–90 kg, and, for active cases, microbiologically confirmed *M. tuberculosis* infection (drug-sensitive or multidrug-resistant). Exclusion criteria were extensively drug-resistant tuberculosis and use of systemic or inhaled immunosuppressive drugs within two weeks prior to sampling.

Participants were consecutively recruited based on specimen availability; none of them objected to the secondary use of their samples and data in the research prior to enrollment, or even afterward. The final cohort comprised 10 participants (4 clinical, 2 subclinical, and 4 undergoing treatment), including males and transgender individuals, but no female participants. De-identified patient-related data are provided in Figure S2A. Limitations included the small study population, variability in residual sputum volume and time to freezing, and low RNA quantity and quality, which further constrained the analysis.

## Strains construction

Oligonucleotides, plasmids and strains are provided in the Supplemental Information. The FdxA merodiploid translational reporter strain was constructed by PCR amplification of the Rv2007c open reading frame together with 200 bp upstream, including the native promoter, followed by in-frame fusion to an oligopeptide linker and the *mCherry* fluorescent marker. The DNA fragments were ligated into the pMV361-derived integrative vector pND200, containing the L5 phage *attB* integration site and a kanamycin resistance cassette, *Eco*RI, *Avr*II, and *Hpa*I restriction sites.

The GroEL2 merodiploid translational reporter strain was constructed by PCR amplification of the Rv0440 open reading frame together with 300 bp upstream, including the native promoter, followed by in-frame fusion to an oligopeptide linker and the *mRuby2* fluorescent marker. The DNA fragments were ligated into a pTTP1A-derived integrative vector containing the Tweety phage integration site and a kanamycin resistance cassette using *Eco*RI and *Spe*I restriction sites. In addition, the GroEL2 reporter strain was co-transformed with the pMV361-derived integrative vector pLP27, containing the L5 phage *attB* integration site and a hygromycin resistance cassette expressing sfGFP from a strong promoter, to serve as an internal normalization control.

An *M. tuberculosis* ATC-inducible knock-down strain was generated using the CRISPRi gene-silencing system. Briefly, the pLJR965-modified L5-integrative plasmid pGM309 was digested with *Bsm*BI and gel-purified. Two complementary single guide RNA (sgRNA) oligonucleotides targeting the non-template strand of *rv2224c* (Hip1) were designed with appropriate overhangs, annealed, and ligated into the *Bsm*BI-digested backbone. Final plasmids were confirmed by sequencing.

All constructs were verified by restriction profiling or sequencing and transformed into *M. tuberculosis* Erdman or H37Rv by electroporation (2.5 kV, 25 µF, 1000 Ω, 2-mm cuvette). Transformants were selected on Middlebrook 7H10 agar containing the appropriate antibiotic.

To generate the His-tagged GroEL2 expression construct, the *M. tuberculosis* Rv0440 open reading frame was cloned into the pET28 expression vector using sequence- and ligase-independent cloning (SLIC). The PCR insert and linearized vector were separately treated with T4 DNA polymerase in the absence of dNTPs to control generate complementary 5′ single-stranded overhangs. The fragments were then mixed to allow annealing, generating a nicked recombinant plasmid that was directly transformed into competent *E. coli* BL21(DE3), where gaps were repaired. The entire plasmid was verified by sequencing.

## Growth curve assay

Exponentially growing primary cultures were diluted to OD_600_ 0.05 in pre-warmed 7H9 complete medium and incubated at 37 °C for 14 days, during which OD_600_ was measured at regular intervals.

## Flow cytometry and cell sorting of *M. tuberculosis* FdxA reporter

Data acquisition and sorting were performed on a Bio-Rad S3e cell sorter using FL1, FL3, and FL4 channels (405, 488, and 561 nm). Data were analyzed with FCS Express 7 (De Novo Software).

For analysis of FdxA_mCherry_ reporter activity, WT *M. tuberculosis* Erdman and FdxA_mCherry_ strains were cultured to exponential phase in complete 7H9 broth or under defined stress conditions. Stationary-phase cultures were incubated for 4 weeks. For fatty acid (FA) condition, 7H9 was replaced with Hartmans–de Bont medium supplemented with 20 mM sodium propionate as the sole carbon source. Hypoxic cultures were incubated in bottles filled to 80% volume for 21 days. Cultures were treated with 0.01% Tyloxapol for 24 h prior to sorting, filtered through 35 µm nylon cell strainer caps in 5 mL round-bottom Falcon tubes, and adjusted to 10^6^ cells/mL. Cells were stained with 3 µM DRAQ7 1 h prior to analysis to exclude permeabilized bacteria.

Singlet and live bacterial populations were defined using forward/side scatter and DRAQ7 exclusion, while doublets and DRAQ7-positive cells were excluded. Initial voltage setting and compensation were calibrated using non-fluorescent or constitutively mCherry-expressing (pGM218) *M. tuberculosis*. FdxA_mCherry_-Dim and -Bright cells were sorted based on fluorescence threshold in purity mode using a 70 µm nozzle. Acquisition rates were typically maintained at 8,000–10,000 events/sec with sorting rates of 6,000–8,000 events/sec (>80% efficiency). Sorted cells were collected into five tubes containing 1 mL of 4x 7H9 or HdB medium. Cell suspensions containing about 5 x 10^6^ events were harvested at 3900 rpm for 25 min at 21 °C, and pellets were stored at −80 °C until RNA isolation.

## RNA extraction from *M. tuberculosis* subpopulations

RNA was extracted from sorted *M. tuberculosis* subpopulations using a hot phenol–Trizol method. Cell pellets were lysed in 400 µL hot lysis buffer (50 mM Tris-HCl pH 8.0, 1 mM EDTA, 1% SDS) and transferred to tubes containing 400 µL hot phenol (pH 6.6) and 400 µL acid-washed glass beads (425–600 µm). Samples were incubated at 65 °C for 15 min with intermittent vortexing, followed by centrifugation at 10,000 rpm for 15 min at 4 °C. The aqueous phase was recovered, mixed with an equal volume of TRIzol, and extracted with 100 µL chloroform per mL of TRIzol. After centrifugation at 10,000 rpm for 10 min at 4 °C, RNA was precipitated from the aqueous phase using 0.5 M LiCl and three volumes of cold 100% ethanol, mixing and incubating for 2 h at −20 °C. RNA was pelleted at 13,500 rpm for 35 min at 4 °C, washed with 80% ethanol, air-dried, and resuspended in 20 µL nuclease-free water. Residual genomic DNA was removed using the TURBO DNA-free kit according to the manufacturer’s instructions, 30 min digestion at 37 °C followed by DNase inactivation and centrifugation. RNA quantity and quality (RNA integrity number, RIN) were determined using the Agilent Bioanalyzer RNA 6000 Pico kit and the Agilent 2100 Bioanalyzer.

## Ultra-low input RNA library preparation and Illumina sequencing

RNAseq libraries were prepared using the SMARTer Stranded Total RNA-Seq Kit v2 – Pico Input (Takara), suitable for 250 pg–10 ng total RNA. The kit employs SMART® (Switching Mechanism At 5′ end of RNA Template) cDNA synthesis technology to generate Illumina-compatible, strand-specific libraries via PCR amplification. Although optimized for mammalian RNA, it can be applied to bacterial RNA using high-quality input. RNA was fragmented prior to first-strand cDNA synthesis to achieve appropriate library sizes.

<u>First-strand cDNA synthesis</u>: 500 pg of RNA samples were fragmented for 3 min for RIN between 5 and 6, or for 4 min for RIN > 7, starting from 1–8 µL of RNA mixed with 1 µL SMART Pico Oligo Mix v2, 4 µL 5x First-Strand buffer, and nuclease-free water up to 7 µL, incubated at 94 °C for either 3 or 4 min, cooled on ice, then first-strand master mix (SMART TSO Mix v2, RNase inhibitor, SMARTScribe Reverse Transcriptase) was added. cDNA synthesis was performed at 42 °C for 90 min, 70 °C for 10 min, then held at 4 °C.

<u>Adapter and index addition</u>: PCR1 master mix, including 2 µL of nuclease-free water, 25 µL of 2x SeqAmp CB PCR buffer, 1 µL of SeqAmp DNA Polymerase, 1 µL of i7 and i5 indexes, was added to 20 µL cDNA. Samples were incubated in a PCR cycler as follows: 94 °C for 1 min; 5 cycles of 94 °C for 15 sec, 55 °C for 15 sec, 68 °C for 30 sec; 68 °C for 2 min; and 4 °C.

<u>Library purification</u>: The partially amplified libraries were purified using AMPure XP beads, washed with 80% ethanol and eluted in nuclease-free water. Each sample was gently mixed with 40 µL of beads and incubated at room temperature for 8 min, then the mixture was kept on a magnetic separator (Takara) for 5 min, and the supernatant was discarded. Samples were washed twice with 80% ethanol, then ethanol was carefully removed. Beads were air-dried for 5 min and resuspended in 22 µL of nuclease free water, incubated at room temperature for 5 min, placed on the magnetic separator for 2 to 4 min, and 20 µL of supernatant were transferred into fresh PCR strips.

<u>Final RNAseq library amplification</u>: PCR2 master mix, including 26 µL of nuclease free water, 50 µL of SeqAmp CB PCR buffer, 2 µL of PCR2 Primers v2, 2 µL of SeqAmp DNA Polymerase, was added to each sample and gently mixed. Samples were incubated in a PCR cycler as follows: 94 °C for 1 min; 15 cycles of 98 °C for 15 sec, 55 °C for 15 sec, 68 °C for 30 sec; 68 °C for 2 min; and 4 °C. <u>Final purification</u>: Amplified libraries were purified using AMPure XP beads, washed twice with 80% ethanol, air-dried for 5 min, and eluted in nuclease-free water. Library concentration was measured using Qubit dsDNA HS kit and validated with the Agilent High Sensitivity DNA kit. Libraries were sequenced in 100-bp paired-end mode (PE100) on an Illumina HiSeq2500 system in rapid run mode.

## Sputum collection and processing

Sputum samples were solubilized and decontaminated using N-acetyl-L-cysteine and sodium hydroxide (NALC–NaOH, NAC-PAC RED®, Alpha-Tec Systems™, Vancouver, USA) according to supplier’s recommendations to preserve mycobacterial viability and reduce contamination from the respiratory tract microbiota. Briefly, available sputum (approximately 5 mL) was mixed with an equal volume of NALC–NaOH, vortexed for 30 sec, and incubated at room temperature for 15 min. The reaction was neutralized with neutralizing buffer (NPC-67®), followed by centrifugation to collect the mycobacteria. The pellet was resuspended in 1 mL of pellet resuspension buffer (PRB™) for routine diagnostic testing, and the remaining volume was stored at −80 °C for subsequent RNA isolation.

## Total RNA isolation from sputum samples

Residual pellets from sputum specimens were resuspended in 500 μL of hot phenol (65°C) and transferred to tubes containing 400 μL of lysis buffer (50 mM Tris-HCl pH 8.0, 1 mM EDTA, 1% SDS) prewarmed at 65°C and 400 μL acid-washed glass beads (425–600 μm). Samples were incubated at 65°C for 15 minutes with intermittent vortexing and centrifuged at 10,000 rpm for 15 min at 4 °C. The aqueous layer was mixed with an equal volume of TRIzol reagent and incubated for 10 min at 4°C. RNA purification was carried out using the Direct-zol RNA MiniPrep kit (Zymo Research). Lysates were mixed with ethanol, applied to Zymo-Spin™ columns, treated with DNase I, washed, and eluted in 25 μL of nuclease-free water and stored −20 °C until quantification and analysis.

## Nanostring analysis of *M. tuberculosis* transcripts ex vivo

Due to low RIN and concentration, 7 µL of total RNA were used with a custom CodeSet targeting mycobacterial genes (Table S3). For hybridization, RNA was combined with reporter and capture probes in hybridization buffer and incubated at 65 °C for 16 h in a thermal cycler. After hybridization, reactions were processed on the nCounter Prep Station using the High Sensitivity protocol and immobilized on the nCounter cartridge. Digital counts were acquired on the nCounter Digital Analyzer, generating a Reporter Code Count (RCC) file per cartridge. Each run included six positive-and eight negative-control probes to determine technical variation and background signal.

## Flow cytometry and cell sorting of *M. tuberculosis* GroEL2 reporter

For analysis of GroEL2 expression, WT H37Rv and GroEL2_mRuby_ or GroEL2_mRuby_-GFP_cyt_ strains were cultured to exponential phase in complete 7H9 broth supplemented with 0.025% Tyloxapol instead of Tween-80. Cultures were vortexed, filtered through the 35 µm strainer, and analyzed using FL-3 or FL-1 and FL-3 channels. Non-fluorescent bacilli and strains constitutively expressing mRuby (pCZ304) or sfGFP (pLP27) were used for initial voltage setting and compensation. Bacteria were gated based on size and background fluorescence, and 80,000 events per sample were collected. GroEL2_mRuby_ reporter cells were sorted into GroEL2_mRuby_-Dim and -Bright (bottom and top 30% of mRuby fluorescence) in purity mode. Approximately 1 x 10^6^ cells were collected into four 5 mL tubes per subpopulation, each containing 1 mL of 4x concentrated 7H9 medium. Sorted fractions were pooled, centrifuged at 3900 rpm for 15 min at 21 °C, and resuspended in 9 mL of complete 7H9. Purity was confirmed by post-sort analysis, and normalization was guided by plating dilutions on 7H10 plates and by event counts during post-sort runs. Normalized subpopulations were used for antibiotic and stress tolerance assays.

## Determination of Minimal Inhibitory Concentration (MIC)

Sorted subpopulations or mid-log phase cultures diluted to an OD_600_ 0.005 were resuspended in complete 7H9 medium without detergent. White-walled 96-well plates were used for ATP-based luminometric measurement of bacterial viability. Wells containing the highest concentration of drug or stressor were filled with 200 µL of bacterial suspension and used to prepare subsequent 2-fold serial dilutions in wells containing 100 µL of bacterial suspension. Plates were incubated under static conditions at 37 °C for 8 days. After incubation, 100 µL of BacTiter-Glo™ reagent (Promega) was added to each well, and plates were incubated for 5 min at room temperature. Luminescence was measured using a Glomax luminometer (Promega) with 0.3 s integration time. Background luminescence from medium-only wells was subtracted, and values were normalized to untreated control wells. Percentage survival was plotted against the logarithm of the drug concentration, and half-maximal inhibitory concentration (IC50) values were determined by fitting a non-linear regression of bacterial viability versus log(concentration) of stressors in GraphPad Prism. For each replicate, IC50 values were compared between bright and dim subsets using an unpaired t-test.

## Macrophage infection

Secondary *M. tuberculosis* cultures were grown to an OD_600_ of 0.5–0.7 prior to infection. Cultures were vigorously vortexed, washed twice with PBS (pH 7.4), and resuspended in complete RPMI 1640 medium at an OD_600_ of 0.5 (approximately 5 x 10^7^ bacilli/mL). M0 THP-1 macrophages were infected at a multiplicity of infection (MOI) of 3 or 5 bacilli per macrophage for 3 h. Extracellular bacteria were removed by washing three times with pre-warmed complete RPMI 1640, and cells were incubated at 37 °C with 5% CO_2_ for either 24 or 48 h prior to downstream assays.

## Lumit immunoassays

THP-1 cells infected with the GroEL2_mRuby_-GFP_cyt_ reporter strain at an MOI of 3 for 24 h were sorted as described above. GroEL2_mRuby_-Bright and -Dim subsets were pelleted, resuspended in fresh RPMI 1640 medium, and 30,000 cells per well were dispensed into white-walled 96-well plates. Cytokine secretion (IL-10, IL-4, TNF-α, IL-1β, and IL-6) was measured using the Lumit Immunoassay™ (Promega) following the manufacturer’s instructions after 18 h incubation at 37 °C, 5% CO_2_. Luminescence was recorded with a Glomax luminometer (Promega) using a 0.3 s integration time. Calibration curves were generated from serial dilutions of individual human cytokines, with medium-only wells used for background subtraction. Cytokine concentrations in samples were interpolated from Log-Log plots of background-subtracted relative luminescence units (RLU) versus cytokine standards. For each replicate, cytokine concentrations were compared between Bright and Dim subpopulations using a paired multiple t-test with Holm-Sidak correction for multiple comparisons.

## Luminescence assay for measurement of reactive oxygen species

THP-1 cells infected with GroEL2_mRuby_-GFP_cyt_ reporter strain at MOI 3 for 48 h were sorted as described above. Bright and dim subsets were pelleted, resuspended in fresh RPMI medium, and 30,000 cells per well were dispensed into white-walled 96-well plates. ROS-Glo H2O2 Assay (Promega) was carried out according to manufacturer’s instructions. Briefly, the H_2_O_2_ substrate (Promega) was added to cells for 2 h at 37 °C, 5% CO_2_, followed by addition of the Ros-Glo Detection solution and a 20-min incubation period at room temperature. Luminescence was recorded using a Glomax luminometer (Promega) with a 0.3 s integration time.

## Annexin V apoptosis imaging assay

THP-1 macrophages were differentiated to an M0 macrophage-like state in CELLview glass-bottom slides (Greiner Bio-One) and infected with the GroEL2_mRuby_-GFP_cyt_ reporter strain at an MOI of 3. At 48 hpi, cells were washed and incubated with 1x annexin-binding buffer (10 mM HEPES, 140 mM NaCl, 2.5 mM CaCl_2_, pH 7.2) containing 5 µL Annexin V–Alexa Fluor 647 conjugate (Thermo Fisher) and 3 µM DRQ7 (Invitrogen) per well for 15 min at room temperature prior to imaging. Images were acquired at 100x magnification, 1024 x 1024 resolution, with seven z-sections of 2 µm each, using the following settings: bright field 10%T, 0.2 sec; FITC (Ex 475/28, Em 525/48 nm), 10%T, 0.1 sec; TRITC (Ex 542/27, Em 597/45), 100%T, 0.2 sec; Cy5 (Ex 632/22, Em 679/34), 100%T, 0.05 sec.

### NF-κB_SEAP and IRF_Lucia reporter THP-1 monocyte assays

WT and KO_MyD88_ THP-1 Dual cells (InvivoGen) were seeded at 40,000 cells per well in 96-well plates, activated with 30 ng/mL PMA for 48 h, and rested in medium alone for 24 h prior to experiments.

For experiments with recombinant GroEL2 (Euromedex), macrophages were pretreated with either 760 nM CLI-095 or 10 µg/mL anti-hTLR2-IgA before addition of GroEL2 protein at varying concentrations. Cells were incubated at 37 °C, 5% CO_2_ for 24 h.

For infection experiments, *M. tuberculosis* strains carrying either the CRISPRi control plasmid (pLP41) or the *rv2224c* (Hip1) CRISPRi plasmid (pLP42) were grown to exponential phase, diluted to OD_600_ 0.05, and cultured with or without 100 ng/mL ATC for 4 days prior to infection. PMA-activated THP-1 Dual reporter cells were pretreated with 760 nM CLI-095 for 1 h, then infected at an MOI of 2 for 2 h, maintaining CLI-095 and ATC where applicable. Cells were washed three times with PBS and incubated at 37 °C, 5% CO_2_ for 24 h before analysis.

NF-κB pathway activity was quantified by secreted embryonic alkaline phosphatase (SEAP) expressed from an IFN-β minimal promoter fused to five NF-κB response elements and three c-Rel binding sites (NF-κB_SEAP). SEAP secretion was measured using the QUANTI-Blue assay (InvivoGen), with 20 µL of cell-free supernatant added to 180 µL of QUANTI-Blue solution, incubated at 37 °C for 1 h, and absorbance read at OD_600_. IRF pathway activity was measured by Lucia luciferase under the ISG54 minimal promoter with five IFN-stimulated response elements (ISRE_Lucia). Lucia luciferase was quantified using the QUANTI-Luc™4 Lucia/Gaussia assay (InvivoGen), adding 50 µL of reagent to 20 µL of cell-free supernatant and recording luminescence immediately at 0.1 s integration time.

## Western blot analysis of GroEL2

WT H37Rv bacteria were grown in 7H9 medium without BSA and stressed with 0.5x MIC of various stressors for 48 h. For experiments quantifying GroEL2 in the supernatant of *rv2224c* (Hip1) CRISPRi and control strains, bacteria were cultured for 4 days in 25 mL of 7H9 in the absence of BSA, with or without 100 ng/mL ATC. Bacteria were pelleted at 3900 rpm for 15 min, and supernatants were collected, with volumes normalized to equivalent bacterial optical density (OD_600_). Supernatants were supplemented with EDTA-free protease inhibitor (cOmplete, Sigma), filtered through 0.22 µm filters, and concentrated using Pierce™ Protein Concentrators PES (3 kDa MWCO) at 3900 rpm to approximately 700 µL (28x concentration depending on the starting culture).

For bacterial pellets, cells were washed once with cold PBS and once with lysis buffer (50 mM Tris-Cl pH 7.5, 50 mM NaCl, 0.5 mM EDTA, 5% glycerol, 1x EDTA-free protease inhibitor), centrifuged at 12000 rpm for 5 min at 4 °C, and resuspended in 300 µL of lysis buffer. Samples were transferred to 2 mL tubes containing 0.5 mm glass beads and lysed using a Precellys Tissue Homogenizer (Bertin Technologies) for three times at 6000 rpm for 1 min. Lysates were centrifuged, and supernatants were decontaminated through 0.2 µm centrifugal microtubes (Pall Life Sciences).

Protein samples were mixed with 4x SDS-PAGE loading dye, heated for 10 min at 95 °C, separated on NuPAGE 4–12% Bis-Tris gels, and transferred to nitrocellulose membranes, using an iBlot 2 system (Invitrogen). Membranes were blocked in PBS + 0.05% Tween-20 (PBST) + 5% non-fat dry milk for 1 h at room temperature, then incubated overnight with primary antibodies: anti-HSP65 (1:2000 in PBST) or anti-Ag85 (1:1000 in TBST). Membranes were washed three times with PBST and incubated for 1 h at room temperature with HRP-conjugated anti-mouse IgG secondary antibody (1:10000 in PBST). Blots were developed using SuperSignal West Atto Ultimate Sensitivity Substrate (Thermo Fisher) and imaged with a Fusion FX system (Vilber).

## Expression and purification of *M. tuberculosis* GroEL2-His

Histidine-tagged GroEL2 protein (GroEL2-His) was purified from *E. coli* BL21(DE3) carrying the pMP2 expression plasmid. Bacteria were precultured overnight in 500 mL LB supplemented with kanamycin (50 µg/mL) at 37 °C and 180 rpm and used to inoculate 2 L culture to an initial OD_600_ of 0.2. Cultures were grown at 37 °C and 120 rpm to OD_600_ 0.8, then expression was induced with 1 mM IPTG and incubated overnight at 20 °C. Cells were harvested by centrifugation at 6000 rpm for 10 min and stored at −80 °C. Pellets were resuspended in lysis buffer (50 mM Tris-HCl pH 7.5, 300 mM NaCl, 10 mM imidazole, cOmplete EDTA-free protease inhibitors (Roche), and Pierce^TM^ universal nuclease for cell lysis (ThermoFisher Scientific) at 5 mL per gram wet weight, lysed using a Cell Disruptor CF1 at 1.3 kBar, and clarified by centrifugation at 55000 × g for 1 h at 4 °C. Cleared lysate was applied to a Protino 5 mL Ni-NTA column (Macherey-Nagel) on an ÄKTA Pure^TM^ chromatography system (Cytiva). The Ni-NTA column was equilibrated with lysis buffer, washed with wash buffer 1 (50 mM Tris-HCl pH 7.5, 300 mM NaCl, 20 mM imidazole) and bound protein was eluted with elution buffer (50 mM Tris-HCl pH 7.5, 300 mM NaCl, 250 mM imidazole). Elution fractions were pooled and dialyzed overnight at 4 °C against 50 mM Tris-HCl pH 8, 300 mM NaCl using a 10 kDa MWCO Slide-A-Lyzer cassette (ThermoFisher Scientific). Dialyzed proteins were concentrated through centrifugation using a 10kDa Amicon® Ultra Centrifugal concentrator (Merck Millipore). Concentrated protein was then loaded onto a HiLoad® 16/600 Superdex® 200 pg gel filtration column equilibrated in 50 mM Tris-HCl pH 8, 300 mM NaCl and eluted at 0.5 mL/min. Fractions with pure GroEL2-His were pooled, concentrated to 3–5 mg/mL, supplemented with 20% glycerol, aliquoted, and stored at −80 °C.

## Endotoxin detection

Endotoxin levels in recombinant GroEL2 were measured using the ToxinSensor™ Chromogenic LAL Endotoxin Assay Kit (GenScript), according to manufacturer’s instructions. Samples were adjusted to pH 6–8 and diluted to fall within the assay linear range (0.01–1 EU/mL). Endotoxin standards were prepared in Limulus Amebocyte Lysate (LAL) reagent reconstituted in water and serially diluted to generate a standard curve. For each assay, 100 µL of sample or standard was mixed with 100 µL of reconstituted LAL reagent, incubated at 37 °C, and then combined with 100 µL of chromogenic substrate solution for a final incubation of 6 min at 37°C. After the reaction, color-stabilizing solutions were added with gentle swirling to avoid foaming. Reactions were transferred to a 96-well plate, and absorbance was measured at 545 nm using a Tecan microplate reader. Sample endotoxin concentrations were calculated from the standard curve. Recombinant HSP65 showed endotoxin levels below the 0.05 EU/mL cutoff.

## Western blot detection of LPS

A range of concentrations of LPS and GroEL2 were mixed with 4x SDS-PAGE loading dye and heated at 95 °C for 5 min prior to loading onto two NuPAGE 4–12% Bis-Tris gels in 1x MES running buffer at 100 V for 90 min. One gel was stained with Coomassie, and the parallel gel was transferred onto PVDF membranes using an iBlot Dry Transfer System (Invitrogen) following the standard 7-min protocol. Membranes were blocked in TBST supplemented with 5% non-fat dry milk overnight at 4 °C, washed five times with TBST, and incubated with primary antibody against *E. coli* O111:B4 LPS at 0.5 µg/mL in TBST containing 1% non-fat dry milk for 24 h at 4 °C. After five washes with TBST, membranes were incubated for 2 h at room temperature with HRP-conjugated goat anti-mouse IgG secondary antibody (1:15000 in TBST with 1% non-fat dry milk). Blots were developed using Pierce ECL Western Blotting Substrate (Thermo Fisher) according to the manufacturer’s instructions and imaged using a Fusion FX system (Vilber).

## GroEL2-His pull-down assay in THP-1 cells

For preparing cell extracts, WT and KO_MyD88_ THP-1 Dual cells (InvivoGen) were seeded at 500,000 cells per well in 12-well plates, activated with 30 ng/mL PMA for 48 h, and rested in medium alone for 24 h prior to stimulation with 100 ng/well of GroEL2-His purified protein. Cells were incubated at 37 °C, 5% CO_2_ for 24 h. After incubation, cells were washed in ice-cold PBS, harvested and resuspended in 200 µL of lysis buffer (50 mM Tris HCl pH 8, 150 mM NaCl, 1% NP40, supplemented with cOmplete protease inhibitors (Roche) and 1 mM PMSF (Sigma). Cells were lysed at 4 °C on a rotating wheel for 1 h and then centrifuged at 20000 x g at 4°C for 10 min to pellet cell debris. Supernatants were transferred to Low-Bind Protein Eppendorf tubes.

For pull-down experiments, 120 µL of Ni-NTA agarose resin (Qiagen) were transferred to a Low-Bind Protein Eppendorf tube, washed three times, and equilibrated in equilibration buffer (20 mM Tris-HCl pH 7.5, 150 mM NaCl). Next, 50 µg of GroEL-His recombinant protein (bait) was mixed with 400 µL of equilibration buffer and incubated with washed beads for 1 h at 4°C on rotation, followed by 10 min on ice in static conditions. Samples were then centrifuged for 1 min at 4 °C at 1000 x g and flow-through was discarded, except for a small fraction kept for further analysis. Similarly, 200 µL of cell supernatants were mixed with equilibration buffer at 1:1 ratio (v/v), loaded on GroEL2-His-bound beads and incubated for 1h at 4 °C on rotation, followed by 10 min on ice in static conditions. Samples were then centrifugated for 1 min at 4 °C at 1000 x g and flow-through was discarded, except for a small fraction kept for further analysis. Beads were washed three times for 5 min on rotation with 400 µL of equilibration buffer. Final wash was supplemented with 20 mM imidazole to decrease non-specific binding. Tubes were then centrifugated for 1 min at 4 °C at 1000 x g and flow-through was discarded, except for a small fraction kept for further analysis. Washed beads were then transferred to a fresh Low-Bind Protein Eppendorf tube, resuspended in 80 µL of elution buffer (20 mM Tris-HCl pH 7.5, 150 mM NaCl, 500 mM imidazole) and further incubated for 10 min at 4 °C on rotation to detach GroEL2-His-interacting proteins. Supernatant was collected by centrifugation at 1000 x g for 1 min at 4°C. Elution step was repeated twice. Fractions were analyzed by SDS-PAGE to validate the pulldown experiment, and elutions were pulled and stored at -80 °C for identification of GroEL2-His-interacting proteins by mass spectrometry.

## Mass spectrometry-based proteomics of GroEL2 pull-down

Protein digestion for mass spectrometry was performed using an adapted version of the S-Trap™ protocol (Zougman et al., 2014) with S-Trap mini columns (ProtiFi™). Briefly, proteins were solubilized in lysis buffer (5% SDS, 50 mM TEAB pH7,55). Cysteines were reduced with 120 mM Tris(2-carboxyéthyl) phosphine hydrochloride, TCEP for 15 min at 55 °C. Then cysteines were alkylated with 500 mM iodoacetamide (IAA) for 10 min at room temperature in the dark. Samples were acidified by adding 12% phosphoric acid at a 1:10 ratio (v/v). S-Trap binding/wash buffer (100 mM AMBIC in 90% methanol/10% water) was added before loading onto the S-Trap column. Columns were centrifuged at 4000 x g for 30 sec to bind proteins to the matrix. Proteins were washed three times with 400 µL of S-Trap binding/wash buffer, each followed by centrifugation at 4000 x g for 30 sec. For enzymatic digestion, sequencing-grade trypsin in 50 mM AMBIC at a 1:10 ratio (enzyme:sample, w/w) was added to each column. After a brief centrifugation to ensure enzyme penetration, digestion was performed at 37 °C in a thermostatic bath overnight. Peptides were sequentially eluted with 50 mM AMBIC, 0.2% formic acid (FA) and 50% acetonitrile (ACN), 0.2% FA. Each elution step was followed by centrifugation at 4000 x g for 30 s. Combined eluates were dried in a SpeedVac concentrator and stored at −20 °C until LC–MS/MS analysis.

For liquid chromatography-mass spectrometry (LC-MS), all peptide fractions were resuspended in 2% acetonitrile (ACN), 0.1% formic acid (FA) immediately before LC–MS/MS injection. Peptide analysis was performed on a timsTOF Ultra mass spectrometer (Bruker) coupled to a nanoElute LC system (Bruker). Peptides were loaded onto a PepSep Twenty-five Series column (75 µm i.d., 1.5 µm particle size, Bruker) and equilibrated in buffer A (0.1% FA) at 800 bar. Chromatographic separation was achieved at a flow rate of 250 nL/min and a column temperature of 50 °C using the following multi-step gradient of buffer B (80% ACN, 0.1% FA): 2–4% in 1 min, 4–20% in 20 min, 20–30% in 5 min, and 30–95% in 1 min. The CaptiveSpray source was operated with a capillary voltage of 1,600 V, a dry gas flow of 3.0 L/min, and a temperature of 200 °C.

MS data were acquired in dia-PASEF (data-independent acquisition parallel accumulation–serial fragmentation) mode. In this acquisition strategy, precursor ions were accumulated in the first TIMS analyzer, separated by ion mobility, and sequentially released into the quadrupole for fragmentation. The full m/z range from 100 to 1,700 Th was covered using optimized isolation windows aligned to ion mobility values spanning 0.64–1.45 Vs/cm² (1/K₀). Each dia-PASEF cycle comprised multiple MS/MS scans with a total duty cycle of ∼1.1 s. Collision energies were ramped linearly as a function of ion mobility, following the manufacturer’s recommendations, to ensure optimal fragmentation across the mobility range.

## Snapshot microscopy

Phase contrast and fluorescence snapshot imaging of *M. tuberculosis* fluorescent reporters under different conditions was carried out using an inverted DeltaVision Elite Microscope (Cytiva) equipped with a UPLFLN100XO2/PH3/1.30 oil objective (Olympus). For each sample, 0.5 µL of culture was placed between two #1.5 coverslips and evenly distributed by gentle rubbing before sealing with glue. The FdxA_mCherry_ reporter in exponential or stationary phase, and after 24 h exposure to 20 mM sodium propionate, 5 mM H_2_O_2_, pH 5.0, 10% human serum, or nutrient starvation in Hartmans-de Bont medium, was imaged under the following conditions: phase contrast 50%T, 0.1 sec; and TRITC (Ex 542/27, Em 597/45), 50%T, 0.2 sec. The GroEL2_mRuby_ reporter in exponential phase and after 48 h exposure to 0.5x MIC of RIF (MIC: 0.038 µg/mL), EMB (MIC: 1.25 µg/mL), MOX (MIC: 0.025 µg/mL), DFO (0.25 mg/mL), NO (0.1 mM), or H_2_O_2_, (2.5 mM) was imaged under the following conditions: phase contrast 100%T, 0.1 sec and TRITC (Ex 542/27, Em 597/45), 100%T, 0.15 sec.

### Time-lapse microscopy of *M. tuberculosis* GroEL2_mRuby_-GFP_cyt_ reporter

Exponential-phase *M. tuberculosis* cultures were diluted 1:10 in complete 7H9 medium and vortexed vigorously to declump. A 25 x 50 mm #1.5 coverslip was placed on a custom metal frame, and the sterile hexa-device (Manina et al., 2019) was positioned on an acrylic layer with channels facing upward. A membrane of equivalent dimensions equal to the hexa-device was prepared from Spectra/Por 6 dialysis tubing (25 kDa, 12mm), rehydrated with pre-warmed 7H9 on ashless Whatman round filter paper, and 2 µL of diluted bacterial suspensions from independent cultures were dispensed onto each of the six observation areas. Bacteria were allowed to settle for 5 min before the membrane was transferred onto the hexa-device. The device was inverted, placed on the coverslip, and secured evenly with eight screws. Two syringes filled with pre-warmed 7H9 were connected to the inlet ports of the hexa-device, and medium was perfused at 10 µL/min using a syringe pump. The assembled device was mounted on the inverted DeltaVision Elite Microscope (Cytiva) equipped with a UPLFLN100XO2/PH3/1.30 oil objective (Olympus). Images were acquired at 3 h intervals using the following exposure conditions: phase contrast, 100%T, 0.1 s; FITC (Ex 492/28, Em 523/23), 50%T, 0.08 s; TRITC (Ex 542/27, Em 597/45), 100%T, 0.2 s. Single-cell analyses were pooled from two independent experiments, each containing three independent cultures on the device.

## Time-lapse microscopy of infected macrophages

THP-1 monocytes were seeded at 30,000 cells per well in 32 µL on 8-well CultureWell™ Chambered Coverglass (Grace Bio-Labs) and differentiated into M0 macrophages by activation with 30 ng/mL PMA for 48 h. Cells were washed and rested in complete RPMI 1640 medium 24 h prior to infection. Exponentially growing GroEL2_mRuby_-GFP_cyt_ reporter bacilli were added at an MOI of 3 for 2 h, followed by three washes with PBS.

Infected macrophages were embedded in a hydrogel layer to provide structural support. The hydrogel was prepared as follows: 8-arm PEG10kDa vinyl-sulfone (PEGVS) was solubilized to 12% (w/v) in triethanolamine (TEA) buffer (0.3 M, pH 7.5), and 4-arm PEG10kDa thiol (PEGTH) was solubilized to 12% (w/v) in deionized H_2_O. A 7% (w/v) PEGVS–PEGTH solution in TEA, with a stoichiometric balance of thiol and vinyl-sulfone groups, was prepared in 3/4 of the final volume. In the remaining 1/4 of the volume, neutralized collagen was prepared at 6 mg/mL and pH 7.5. The PEGVS–PEGTH solution and collagen were mixed, and 18.3 µL of hydrogel was added per well.

The coverslip was then transferred onto the hexa-device (Manina et al., 2019) so that the wells were in contact with the serpentine channels. The device and the chambered coverglass were inverted, placed on a custom-made metal frame and secured with 8 screws (Figures S5F and S5G). Two syringes filled with pre-warmed RPMI 1640 were connected to the inlet ports of the hexa-device, and medium was perfused at 1.8 µL/min using a syringe pump. The assembled device was mounted on the inverted DeltaVision Elite microscope (Cytiva) equipped with a PlanApo N 60X/1.42 oil objective (Olympus). Images were acquired at 4 h intervals using a 1024 x 1024 pixels field of view and an sCMOS camera, with seven z-sections spanning 12.8 µm combined by maximum intensity projection. Exposure conditions: phase contrast, 32%T, 0.08 sec; FITC (Ex 492/28, Em 523/23), 10%T, 0.05 sec; TRITC (Ex 542/27, Em 597/45), 100%T, 0.1 sec. Experiment was repeated independently at least three times.

## Synthesis of GroEL2 inhibitor mr38548

<u>Preparation of 4-Nitro-(2-aminothioxomethyl)benzoic acid hydrazide</u>: a solution of 4-nitrobenzoic acid (1.1 eq., 1.1 mmol) in SOCl2 (2 mL) under N2 was heated at 80 °C for 1 h. Solvent was evaporated and the residue was solubilized in CH3CN (2 mL). This solution was added dropwise to a mixture of thiosemicarbazide (1.0 eq., 1.0 mmol) and pyridine (3 droplets) in CH3CN (3 mL) at 0 °C. The mixture was stirred at rt for 2 h. The precipitate was filtered, rinsed with CH3CN and recrystallized in EtOH to afford the pure product with a 68% yield. ^1^H NMR (600 MHz, DMSO-d_6_) δ = 10.71 (s, 1H), 9.44 (s, 1H), 8.33 (d, *J* = 8.8 Hz, 2H), 8.12 (d, *J* = 8.8 Hz, 2H), 7.96 (s, 1H), 7.79 (s, 1H). LC-MS (M+H^+^): m/z 241.25 (Sarkar et al., 2022).

<u>Preparation of 3-(4-Nitrophenyl)-5-mercapto-4H-1,2,4-triazole</u>: to a solution of 4-Nitro-(2-aminothioxomethyl)benzoic acid hydrazide (1.0 eq., 1 mmol) in water (8 mL) were added NaOH pellets (10 eq., 10 mmol). The solution was stirred at 20 °C for 1 h. The mixture was acidified with an aqueous solution of HCl 3N and the precipitate was filtered and rinsed with H_2_O to afford the desired product with 50% yield. ^1^H NMR (600 MHz, DMSO-d_6_) δ = 14.18 (s, 1H), 13.95 (s, 1H), 8.37 (d, *J* = 8.9 Hz, 2H), 8.16 (d, *J* = 8.9 Hz, 2H). LC-MS (M+H^+^): m/z 223.23 (Sarkar et al., 2022).

<u>Preparation of 3-(4-Nitrophenyl)-5-(prop-2-ynylthio)-*4H*-1,2,4-triazole (mr38458)</u>: to a solution of 3-(4-Nitrophenyl)-5-mercapto-4H-1,2,4-triazole (1.0 eq., 1 mmol) in MeOH (4 mL) was added EtONa (10 eq., 10 mmol). The solution was stirred at rt for 15 min. Then, propargyl bromide (1.0 eq., 1 mmol) was added dropwise at 0 °C. The mixture was stirred at rt for 30 min. The mixture was poured into ice, and the precipitate was filtered, rinsed with MeOH and recrystallized in EtOH to afford the desired product with 48% yield. ^1^H NMR (600 MHz, Chloroform-*d*) δ = 8.29 (d, *J* = 8.9 Hz, 2H), 8.24 (d, *J* = 8.9 Hz, 2H), 3.94 (d, *J* = 3.5 Hz, 2H), 2.35 (t, *J* = 3.5 Hz, 1H). LC-MS (M+H^+^): m/z 261.28 (Sarkar et al., 2022).

## Microscale thermophoresis

The Monolith His-Tag Labeling Kit RED-tris-NTA 2nd Generation (NanoTemper) was used to assess the affinity of the dye for the target GroEL2 protein according to the manufacturer’s instructions. Briefly, a 5 µM stock solution of RED-tris-NTA 2nd generation dye was prepared in assay buffer (PBS-T 0.01%, 300 mM NaCl, pH 7.4) and diluted to 50 nM. A 4 µM solution of His-tagged GroEL2 protein was freshly prepared in the same assay buffer. In a 96-well plate, a serial dilution of the His-tagged GroEL2 protein was prepared using 10 µL per well. Then, 10 µL of the diluted dye were added to each well, mixed by pipetting up and down, and incubated for 30 min at room temperature. A Monolith capillary was then dipped into each well, and the assay was performed using a Monolith NT.115 instrument (NanoTemper).

For protein labeling (K_d_ > 10 nM), 100 µL of a 100 nM dye solution and a 20 nM solution of labeled protein were prepared and mixed at a 1:1 ratio (v/v). The mixture was incubated for 30 min at room temperature. The sample was then centrifuged at 15000 x g for 10 min, and the supernatant was transferred to a fresh tube. Three Monolith capillaries were dipped into the labeled protein sample, and three additional capillaries were dipped into assay buffer only as controls. The assay was then performed to verify protein detection.

For the binding check, a 40 nM solution of labeled GroEL2 protein was mixed at a 1:1 ratio (v/v) with a ten-fold diluted ligand (mr38548) in assay buffer. A ten-fold diluted ligand buffer was also prepared. For the target-only sample, 20 μL of 40 nM labeled protein was mixed with 20 μL of diluted ligand buffer. For the complex sample, 20 μL of 40 nM labeled protein was mixed with 20 μL of diluted ligand. For the control sample, 20 μL of diluted ligand was mixed with 20 μL of assay buffer. Three Monolith capillaries were dipped into the target-only, complex, and control samples, respectively, and the assay was performed.

For binding affinity analysis, 200 μL of 40 nM labeled protein solution, 30 μL of ten-fold diluted ligand in assay buffer, and 200 μL of ten-fold diluted ligand buffer were prepared. In a 96-well plate, a serial dilution of the ligand was prepared using 10 μL per well. Subsequently, 10 μL of labeled protein was added to each well and mixed. A Monolith capillary was then dipped into each well, and the assay was performed.

## Quantification and statistical analysis

### Single-cell image analysis

Snapshot microscopy images were analyzed by automated segmentation of single *M. tuberculosis* cells in phase contrast using a customized version of Omnipose, trained on mycobacterial images, coupled with a Python notebook that extracts single-cell size and fluorescence and subtracts background fluorescence from the inverted masks (Mistretta et al., 2024). Time-lapse microscopy videos were analyzed using Napari-Omnibacteria, a customized framework that integrates semi-automated segmentation and temporal tracking with interactive correction in a Napari interface (https://gitlab.pasteur.fr/iah-public/napari-omnibacteria). Manual curation was performed on both segmentation and tracks, rebuilding labels frame by frame and recomputing single-cell trajectories after correction.

Single-cell growth rates were calculated by fitting cell size measurements over the generation time to an exponential (Malthusian) growth model in GraphPad Prism. The resulting growth rate for each cell was then plotted against the background-subtracted mean GroEL2_mRuby_ fluorescence measured over the lifetime of the cell.

For infection snapshots, intracellular bacterial foci were manually segmented using the Selection Brush Tool in Fiji (Schneider et al., 2012) to obtain fluorescence intensity and size measurements. Mean intensity values for both the GroEL2_mRuby_ reporter and the constitutive GFP_cyt_ reporter were background-subtracted, and the ratio between the two was calculated. For time-lapse videos of infection, background-subtracted intensities for each channel were measured at each time point over the lifetime of individual macrophages. The intensity of each channel was averaged across time, and the GroEL2_mRuby_ to GFP_cyt_ ratio was calculated. The area occupied by bacilli for each infectious focus within a macrophage was measured, and growth rates were determined by fitting an exponential (Malthusian) growth model to single-focus size measurements using GraphPad Prism.

## Subpopulation transcriptomics analysis

RNA-seq datasets were produced from subpopulations sorted during three independent experiments. The RNA-seq pipeline from Institut Curie (v0.2.4) was used to perform data analysis. Briefly, reads were trimmed from adapters and mapped to *M. tuberculosis* Erdman genome using the STAR mapper (v2.5.3a). The STAR option ’--quantMode GeneCounts’ was used to produce the count matrix from the gene annotation of *M. tuberculosis* Erdman (ATCC 35801) genome (ASM35020v1.41), assigning reads to features with strand-specific information. Quality control statistics were reported using MultiQC 1.5 (Ewels et al., 2016). Features (genes) differentially expressed between subpopulations (Bright and Dim) were obtained for each culture condition. A standard RNAseq differential analysis was performed, including data normalization, quality assessment of the dataset by graphical exploration of raw and normalized data, test for differential expression for each feature between the conditions, raw P-value adjustment. This analysis was performed using the R software (The R Project for Statistical Computing), Bioconductor (Gentleman et al., 2004) packages including DESeq2 (Anders et al., 2010; Love et al., 2014) and the SARTools package (Varet et al., 2016) developed at Institut Pasteur. Normalization and differential analysis were carried out according to the DESeq2 model and package.

### Analysis of *M. tuberculosis* transcripts from sputum

RNA extracted from sputum samples was analyzed on the nCounter system (NanoString, Bruker) using a custom-designed probe codeset (Table S3). Of all samples, only ten were retained for analysis, as 39–99% of probes were detected above the negative control threshold; the remaining samples showed no detectable signal. Analysis was performed on nSolver Analysis Software (version 4.0). Raw counts were background-subtracted using the mean of eight negative-control probes, normalized to the geometric mean of six positive controls to correct for technical variation, and further normalized to the geometric mean of three housekeeping genes (*gyrA*, *rpoZ*, and *sigA*), to account for differences in the RNA input. Comparisons between treated and clinical tuberculosis patients, or between subclinical and clinical tuberculosis patients, were made using a heteroscedastic Welch’s t-test (P < 0.05), with ratios, log_2_ fold-changes, and uncorrected P values calculated. Unsupervised hierarchical clustering analysis was performed in GraphPad Prism 10 using standardized values (mean = 0, SD = 1), Euclidean distance, and complete linkage.

## MS-based proteomics analysis

For protein identification and quantification, peptide masses were searched using Spectronaut ver. 20.3 software against a UniProt *Homo sapiens* database containing the canonical and reviewed protein sequences (83,374 entries the 2025/04/24) to which the GroEL2-His sequence was added.

Digestion enzyme specificity was set to Trypsin/P and semi specific with up to two missed cleavages allowed. Search criteria included carbamidomethylation of cysteine as a fixed modification, oxidation of methionine and acetylation (protein N terminus) as variable modifications. The minimum peptide length was set to 7 amino acids, and a false discovery rate (FDR) cutoff of 1% was applied at the peptide and protein levels. Otherwise, default settings were used.

For pairwise comparison, protein intensities were normalized by condition using the median-centering function from DAPAR R package (Wieczorek et al., 2017), and their missing values were imputed using the impute.mle function from the imp4p R package (Giai Gianetto et al., 2020). This algorithm imputes values in a condition only when intensity has been quantified in at least one of the samples from the considered condition. To determine if a protein is significantly differentially abundant between two conditions, a moderated *t*-test was performed using the limma R package (Ritchie et al., 2015). Moreover, peptides quantified in one condition but not in the other were also considered differentially abundant. An adaptive Benjamini-Hochberg procedure was applied to the resulting P values using the adjust.p function from the R package cp4p (Giai Gianetto et al., 2016) and a 5% FDR threshold was used to select proteins evolving differently between conditions.

## Statistics

Statistical analyses and data visualization were performed using GraphPad Prism 10, unless otherwise specified. Plots combine data from at least two independent biological replicates, and statistical analyses were computed with n ≥ 3. Statistical tests, sample sizes, and P values are indicated in the corresponding figures or figure legends. Parametric or nonparametric tests were applied depending on whether the data passed normality testing. The alpha risk for the statistical tests was set to 5%.

---

**Figure S1.**
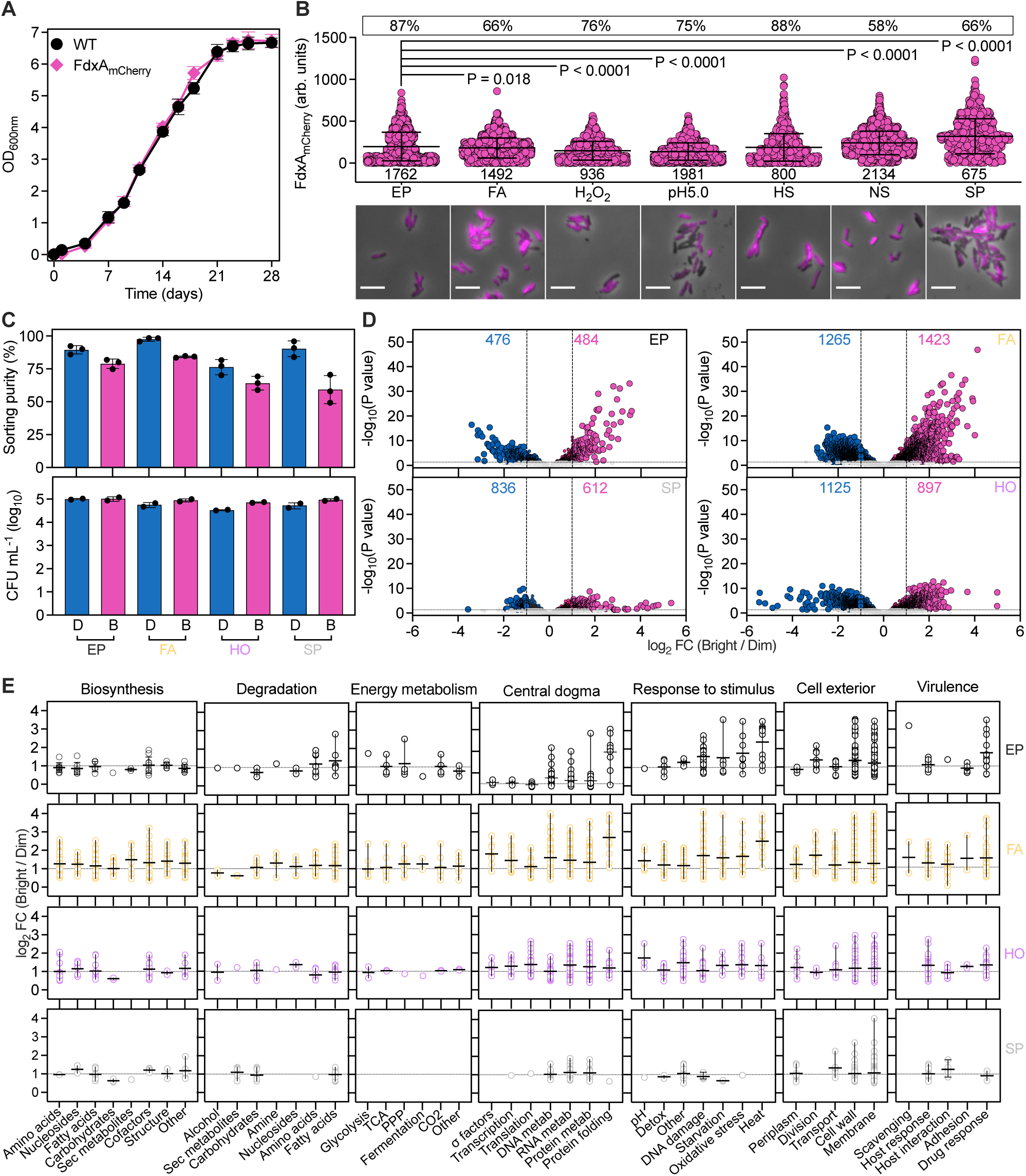
**Single-cell and subpopulation analyses of FdxA_mCherry_ reporter** (A) Growth kinetics of *M. tuberculosis* wild type (WT, black circles) and FdxA_mCherry_ reporter (pink diamonds). Symbols represent mean ± SD (n = 3). (B) Single-cell analysis and representative snapshot images of FdxA_mCherry_ reporter at 4-day exponential phase (EP) or 4-week stationary phase (SP) of growth, and upon 24-hour exposure to: sodium propionate (20 mM) as a short-chain fatty-acid (FA) source; 5 mM H_2_O_2_; low pH (pH5.0); 10% human serum (HS); and nutrient starvation (NS) in Hartmans-de Bont minimal medium. Total number of bacilli is shown at the bottom of the graph and lines represent mean ± SD from 2 independent experiments. Percentage coefficient of variation is boxed. Significance by Brown-Forsythe and Welch ANOVA test (P < 0.0001), followed by Games-Howell’s multiple comparisons test (adjusted P values). Phase contrast (gray) and FdxA_mCherry_ (magenta) are merged. Scale bars, 5 μm. (C) Post-sorting purity from 3 independent experiments and cell density (n = 2) of FdxA_mCherry_-Dim (D) versus -Bright (B) subpopulations in EP, FA, hypoxia (HO), and SP conditions. Lines represent mean ± SD. (D) Numbers of DEGs in FdxA_mCherry_-Bright (pink) versus -Dim (blue) subpopulations, shown in volcano plots ranked by –log_10_ adjusted P values (Wald test with Benjamini-Hochberg correction) and log_2_ fold-changes (FC). Dashed lines mark the ± 2 FC threshold. Environmental conditions as in (C). Dotted line represents the significance threshold (P < 0.05). (E) BioCyc pathway enrichment analysis (Karp et al., 2019; Paley et al., 2017) of significant DEG (P < 0.05) upregulated in FdxA_mCherry_-Bright versus -Dim subpopulations under different conditions as in (D), against *M. tuberculosis* H37Rv database (NCBI 83332, version 28.0, https://mycobacterium.biocyc.org/?sid=biocyc17-3903587489).

**Figure S2.**
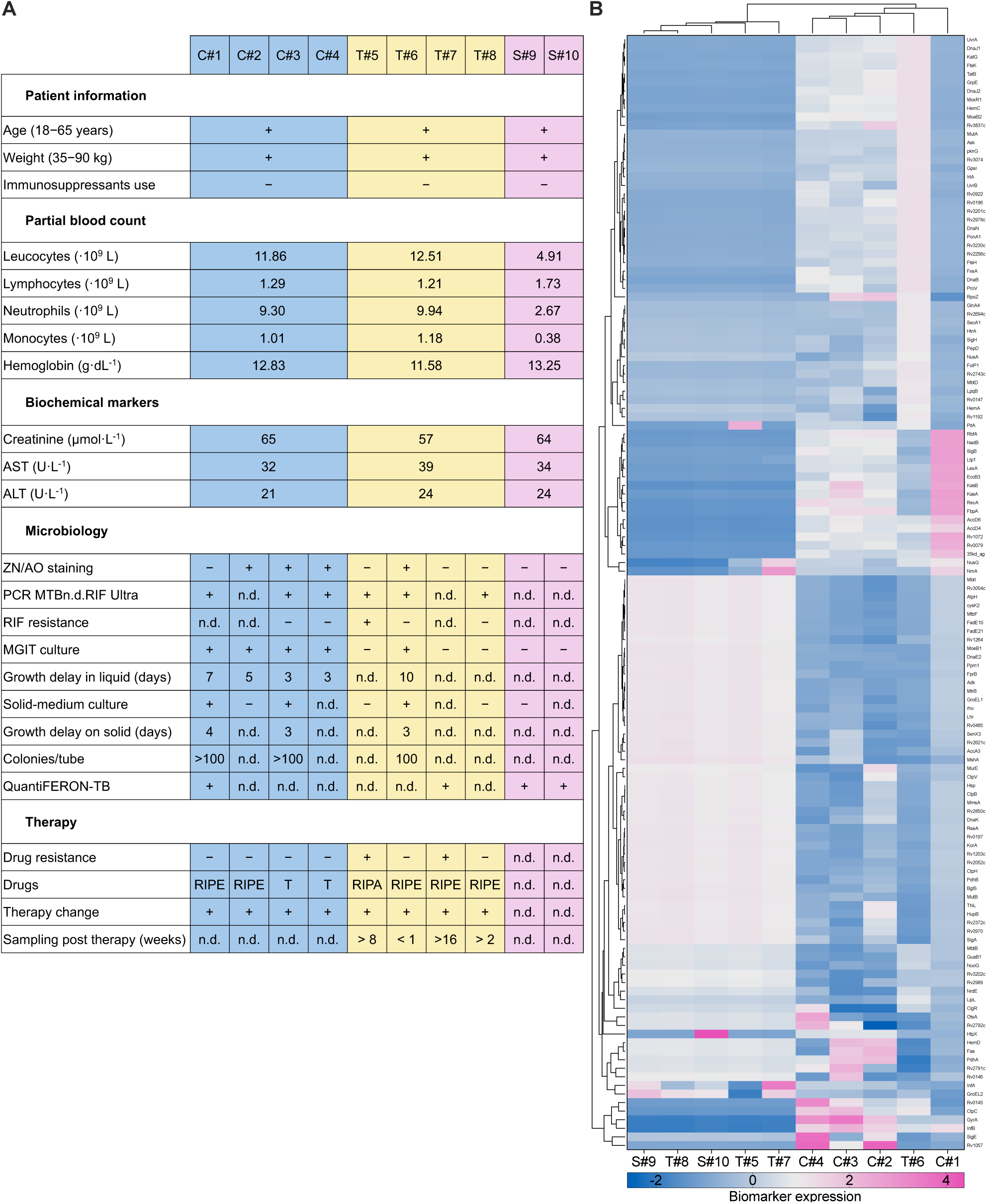
Clinical data and *M. tuberculosis* gene expression profiles from sputum **(A)** Summary of clinical, biochemical, and microbiological information for ten de-identified patients with clinical (C), subclinical (S) tuberculosis, or during different stages of treatment (T). Partial blood count parameters and biochemical markers are reported as averages per patient group. Microbiology: Ziehl-Neelsen and Auramine O (ZN/AO) staining method; Xpert molecular test for rifampicin resistance (PCR MTB/RIF Ultra); mycobacteria growth indicator tube (MGIT); interferon-gamma release assay (QuantiFERON-TB). Drugs: rifampicin (RIF or R); isoniazid (I); pyrazinamide (P); ethambutol (E); tedizolid (T); aminoglycoside (A); fluoroquinolone (F). Parameters are reported as numerical values, positive (+) or negative (-), or not determined (n.d.). **(B)** Unsupervised hierarchical clustering of normalized gene expression data for 130 *M. tuberculosis* stress-response biomarkers detected in sputum samples, measured with the nCounter Analysis System and processed in nSolver 4.0 and GraphPad Prism 10. The heatmap shows expression values, with dendrograms for patients (x-axis) as in panel (A) and genes (y-axis).

**Figure S3.**
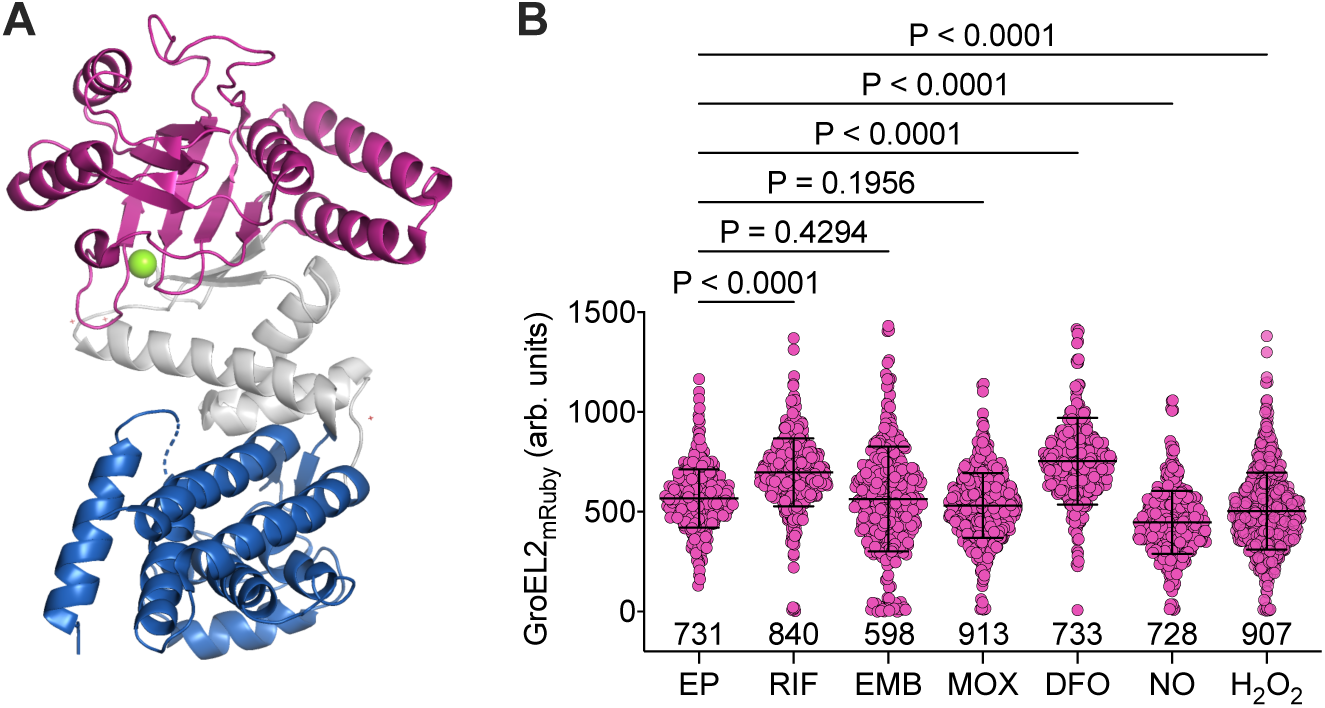
*M. tuberculosis* GroEL2 at the molecular and single-cell level **(A)** Orthoscopic view of the 2.80-Å three-dimensional structure of GroEL2 (Cpn60.2) monomer from *M. tuberculosis* (DOI: https://doi.org/10.2210/pdb3RTK/pdb; Shahar et al., 2011). The three main domains of the 3RTK PDB structure were recolored in PyMOL: apical (residues 190-373, pink); intermediate (residues 134-189 and 374-407, gray); equatorial (residues 1-133 and 408-540, blue). The first 61 and the last 26 residues were not resolved. The magnesium ion ligand is shown in green. **(B)** Single-cell snapshot microscopy analysis of *M. tuberculosis* GroEL2_mRuby_ reporter during exponential phase (EP) of growth and after 48-hour exposure to subinhibitory concentrations of antimicrobials or host-mimetic stressors: rifampicin (RIF, MIC: 0.038 µg/mL), ethambutol (EMB, MIC: 1.25 µg/mL), moxifloxacin (MOX, MIC: 0.025 µg/mL), deferoxamine (DFO, 0.25 mg/mL), nitric oxide (NO, 0.1 mM), and hydrogen peroxide (H_2_O_2_, 2.5 mM). The number of bacilli is shown at the bottom of the graph. Lines represent mean ± SD from 2 independent experiments. Statistical significance by two-way ANOVA (P < 0.0001), followed by with Dunnett’s multiple comparisons test (adjusted P values).

**Figure S4.**
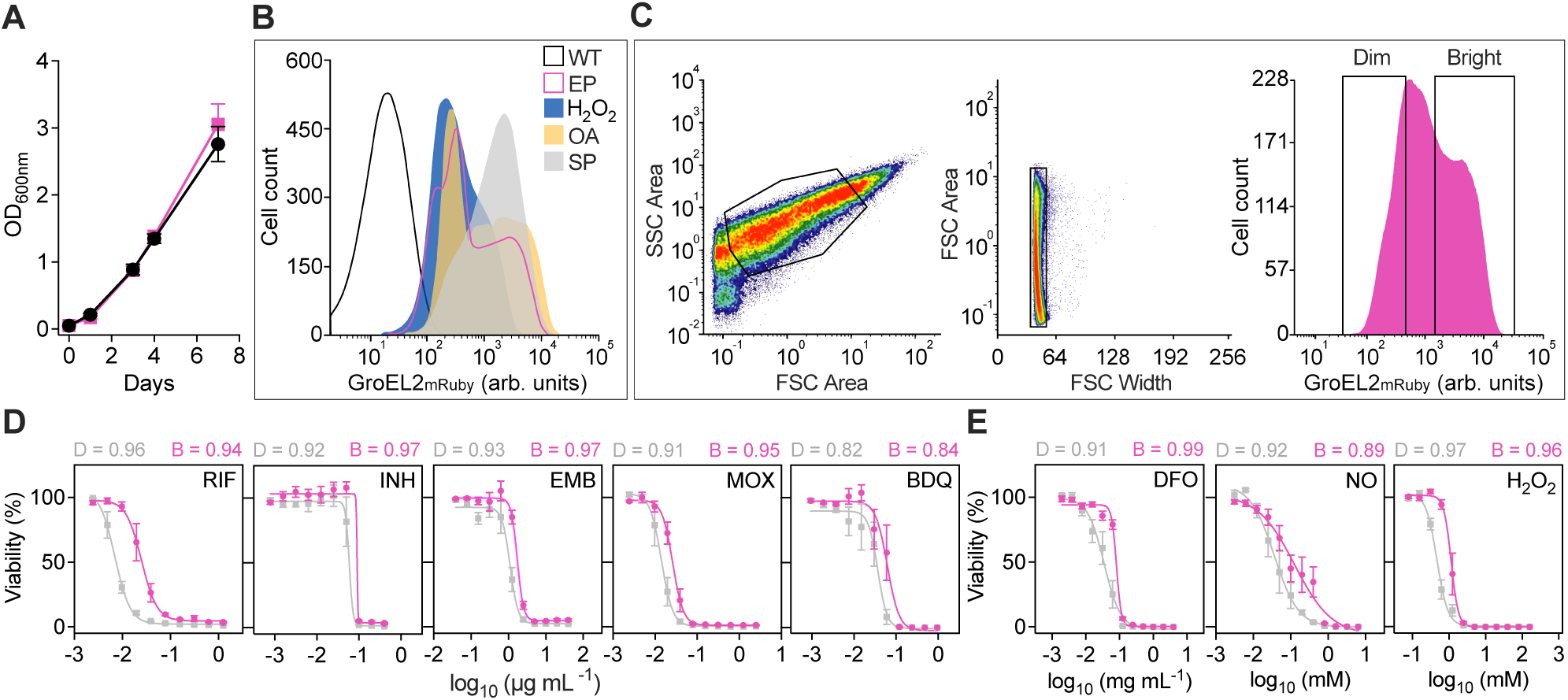
Characterization of GroEL2_mRuby_ *M. tuberculosis* subpopulations **(A)** Growth kinetics of exponentially growing *M. tuberculosis* WT (circle) and GroEL2_mRuby_ reporter (square). Lines represent mean ± SD (n = 3). **(B)** Flow cytometry analysis of WT *M. tuberculosis* (white, black outline) and GroEL2_mRuby_ reporter in exponential phase (EP, pink outline) or stationary growth-phase (SP, gray), and after 48-hour exposure to 5 mM H_2_O_2_ (blue) or 200 µM oleic acid (OA, yellow). Representative overlays are normalized to 100,000 events (n ≥ 3). **(C)** Gating strategy for GroEL2_mRuby_ reporter in EP. Bacterial singlets gating (black polygons). GroEL2_mRuby_-Dim and -Bright subpopulations were gated in the resulting histogram prior to sorting. **(D and E)** Dose-response curves relating concentrations of anti-tubercular drugs (D) and host-mimetic stressors (E) to the viability of GroEL2_mRuby_-Dim (D, gray squares) and -Bright (B, pink circles) subpopulations. Lines represent mean ± SD of at least 4 independent experiments. Data are fit to a four-parameter sigmoidal nonlinear regression model with log-transformed concentrations. Goodness of fit (R^2^) is shown on top of the graphs for each subpopulation.

**Figure S5.**
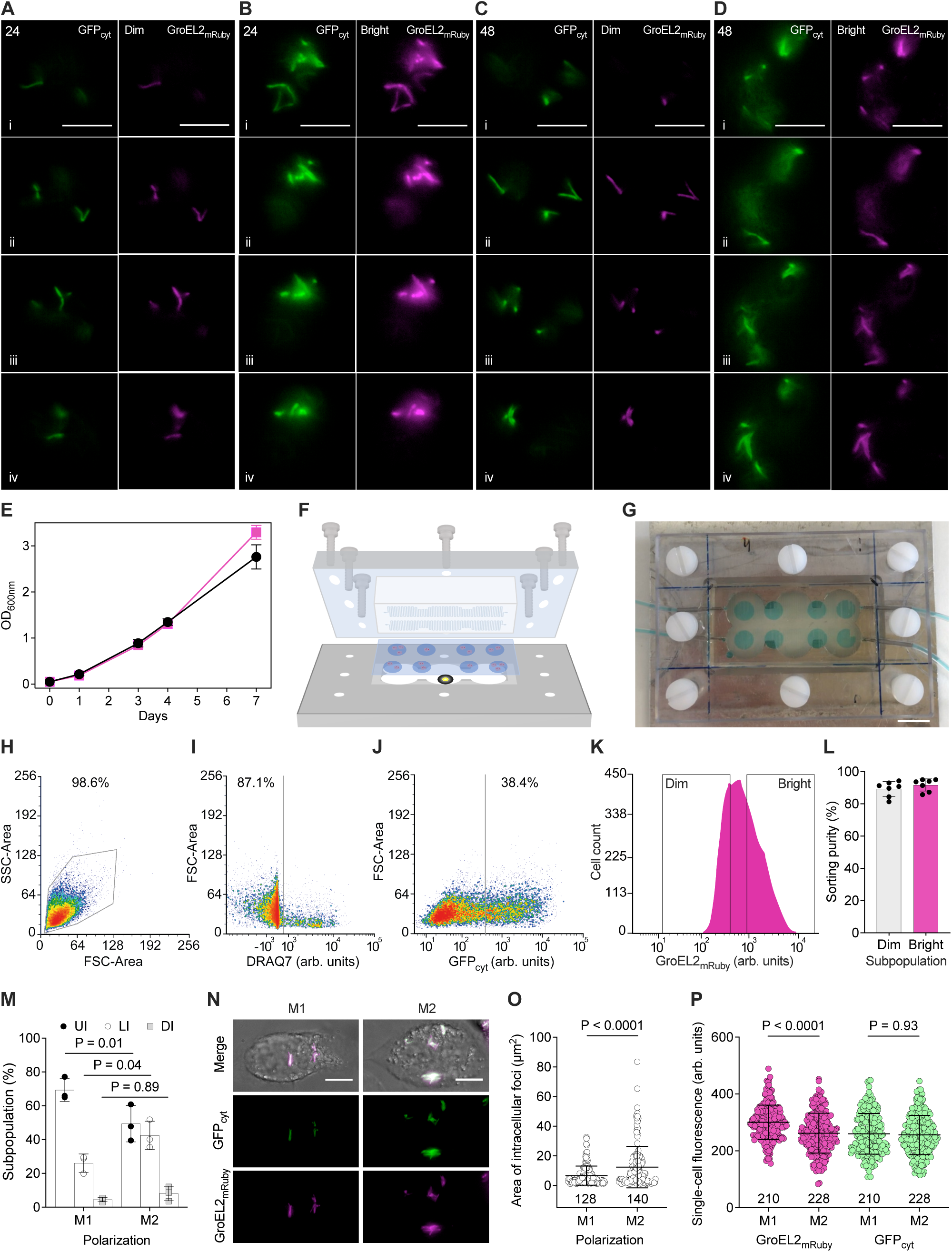
*M. tuberculosis* GroEL2_mRuby_-GFP_cyt_ in resting and polarized THP-1 macrophages (A–D) THP-1 cells infected with GroEL2_mRuby_-GFP_cyt_ reporter at 24 and 48 hpi. Individual z-sections (i to iv) from dashed boxes within maximum intensity projections in Figures 5A–5D. GroEL2_mRuby_-Dim or -Bright (magenta) and GFP_cyt_ (green) channels are shown separately. Scale bars, 10 µm. **(E)** Growth kinetics of exponentially growing *M. tuberculosis* WT (circles) and GroEL2_mRuby_-GFP_cyt_ reporter (squares). Lines represent mean ± SD (n = 3 independent experiments). **(F)** Perspective view schematic of the microfluidic setup used for time-lapse microscopy of infected macrophages seeded on a 6-mm culture-well chambered coverslip (24 × 50 mm) and filled with hydrogel. A polydimethylsiloxane microfluidic device (Manina et al., 2019) was placed on the coverslip, with two serpentine microchannels allowing continuous perfusion of medium via inlet and outlet ports connected to a syringe pump. The assembly is stabilized using a custom-made acrylic and metal frame and screws and mounted on the microscope stage for multi-point time-lapse inverted microscopy. **(G)** Picture of the microfluidic setup perfused with blue dye to visualize the culture-well chambers. Scale bar, 10 mm. **(H–K)** Representative gating strategy for resting THP-1 macrophages infected with GroEL2_mRuby_-GFP_cyt_ reporter at 24 hpi (n = 7 independent experiments). Forward versus side scatters (FSC_ vs SSC_Area) gating (H, black polygon). Live infected cells were selected (black lines) by DRAQ7 negative staining (I) and GFP_cyt_ positive fluorescence (J). GroEL2_mRuby_-Dim- and -Bright subpopulations were gated in the resulting histogram (K). (L) Post-sorting purity of THP-1 macrophages carrying GroEL2_mRuby_-Dim and -Bright subpopulations at 24 hpi. Lines represent mean ± SD (n = 7 independent experiments). (M) Infection rate in pre-polarized M1 or M2 THP-1 macrophages infected with GroEL2_mRuby_-GFP_cyt_ reporter at 24 hpi. DRAQ7-negative staining was used to determine THP-1 viability. Percentages of cells that were uninfected (UI, black circles), live infected (LI, empty circles), and dead infected (DI, gray squares) are shown. Lines represent mean ± SD (n = 3 independent experiments). Significance by two-way ANOVA with Sidak correction for multiple comparisons. (N) Representative snapshot images of live infected THP-1 macrophages pre-polarized into M1 or M2 states at 24 hpi. Maximum intensity projections of six images from a z-stack of 12 µm. Bright field (gray), GFP_cyt_ (green), and GroEL2_mRuby_ (magenta) channels are merged, and fluorescence channels are also shown separately. Scale bars, 10 µM. (O) Size of *M. tuberculosis* GroEL2_mRuby_-GFP_cyt_ infectious foci in pre-polarized THP-1 macrophages. Sample number is shown at the bottom of the graphs. Lines represent mean ± SD (n = 3 independent experiments). Significance by two-tailed unpaired *t*-test (95% confidence level). (P) GroEL2_mRuby_ and GFP_cyt_ fluorescence of intracellular bacilli in pre-polarized THP-1 macrophages 24 hpi. Lines represent mean ± SD (n = 3, with number of infectious foci shown at the bottom of the graphs). Significance by one-way ANOVA (P < 0.0001), followed by Tukey multiple comparisons test (adjusted P values).

**Figure S6.**
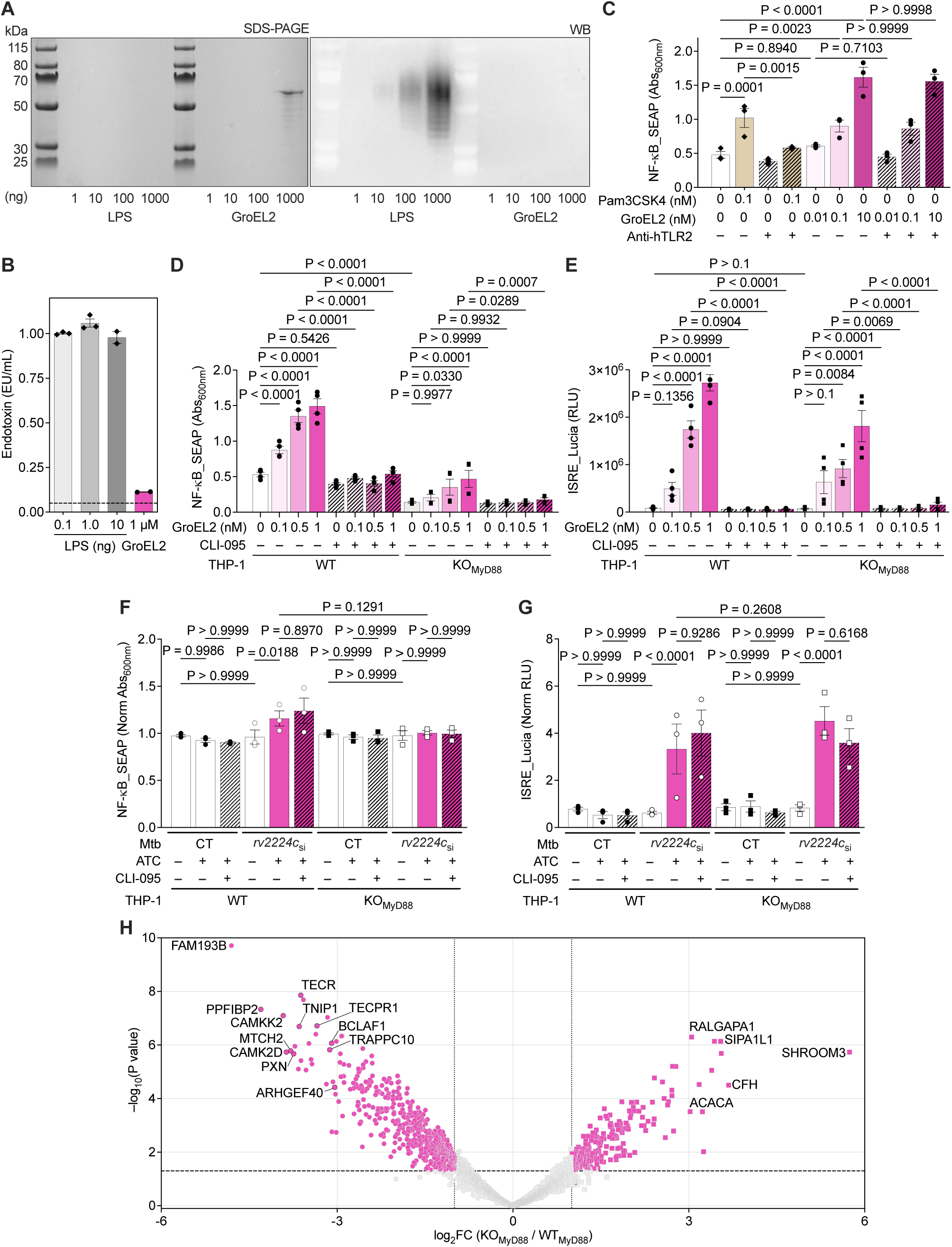
GroEL2 triggers TLR4-mediated inflammatory signaling **(A)** Coomassie-stained SDS-PAGE (left) and immunoblotting with anti-LPS mouse monoclonal IgM (right) of increasing amounts of *E. coli* LPS (Sigma) and recombinant HSP65 (GroEL2) recombinant protein (Euromedex). LPS was not detected in the GroEL2 protein. **(B)** Quantification of endotoxin units (EU) in *E. coli* LPS and recombinant GroEL2 protein at 1 µM. The dashed line represents the threshold for endotoxin contamination. Data are representative of at least two independent experiments. The maximum concentration of GroEL2 used to stimulate THP-1 dual-reporter cells was 1000-fold lower (1 nM), excluding the risk of non-specific TLR4 induction. **(C)** Activation of NF-κB in WT THP-1 dual-reporter cells stimulated with different concentrations of TLR1/2 agonist Pam3CSK4 (diamonds) or GroEL2 recombinant protein (circles) for 24 hours. Absence (–) or presence (+) of anti-hTLR2 mouse monoclonal IgA2 (10 µg/mL) is indicated. Significance by one-way ANOVA for matched datasets (P < 0.0001), followed by Šidák’s multiple comparisons test (adjusted P values). Lines represent mean ± SEM (n = 3). **(D and E)** Activation of NF-κB (D) and IRF (E) pathways in WT (circles) and KO_MyD88_ (squares) THP-1 dual-reporter cells stimulated with different concentrations of GroEL2 recombinant protein for 24 hours. Absence (–) or presence (+) of TLR4 inhibitor CLI-095 (760 nM) is indicated. Significance by one-way ANOVA (P < 0.0001), followed by Šidák’s multiple comparisons test (adjusted P values). Lines represent mean ± SEM (n = 4). **(F and G)** Activation of NF-κB (F) and IRF (G) pathways in WT (circles) and KO_MyD88_ (squares) THP-1 dual-reporter cells 6 hpi. Cells were infected with *M. tuberculosis* CT (filled symbols) or *rv2224c*_si_ (open symbols) at MOI 2 for 2 hours, 4 days after induction of silencing. Absence (–) or presence (+) of inducer and TLR4 inhibitor (760 nM) are indicated. Values are normalized to uninfected controls. Significance by one-way ANOVA for matched datasets (P < 0.0001), followed by Šidák’s multiple comparisons test (adjusted P values). Lines represent mean ± SEM (n = 3). **(H)** Volcano plot of differentially expressed proteins in KO_MyD88_ versus WT_MyD88_ THP-1 cells, upon co-immunoprecipitation with *M. tuberculosis* GroEL2-His (n = 5 independent replicates). Proteins are ranked in a volcano plot based on their –log_10_(P value) derived from a LIMMA t-test (y-axis) versus log_2_-transformed abundance ratio between KO_MyD88_ (squares) and WT_MyD88_ cells (circles). Dotted lines indicate a 2-fold chance (log_2_ FC±1). The dashed line indicates a 1% false discovery rate (FDR). Proteins with FDR <0.01 are considered significantly more abundant in one condition than the other. Identifiers of functionally relevant proteins are indicated (Table S4).

**Figure S7.**
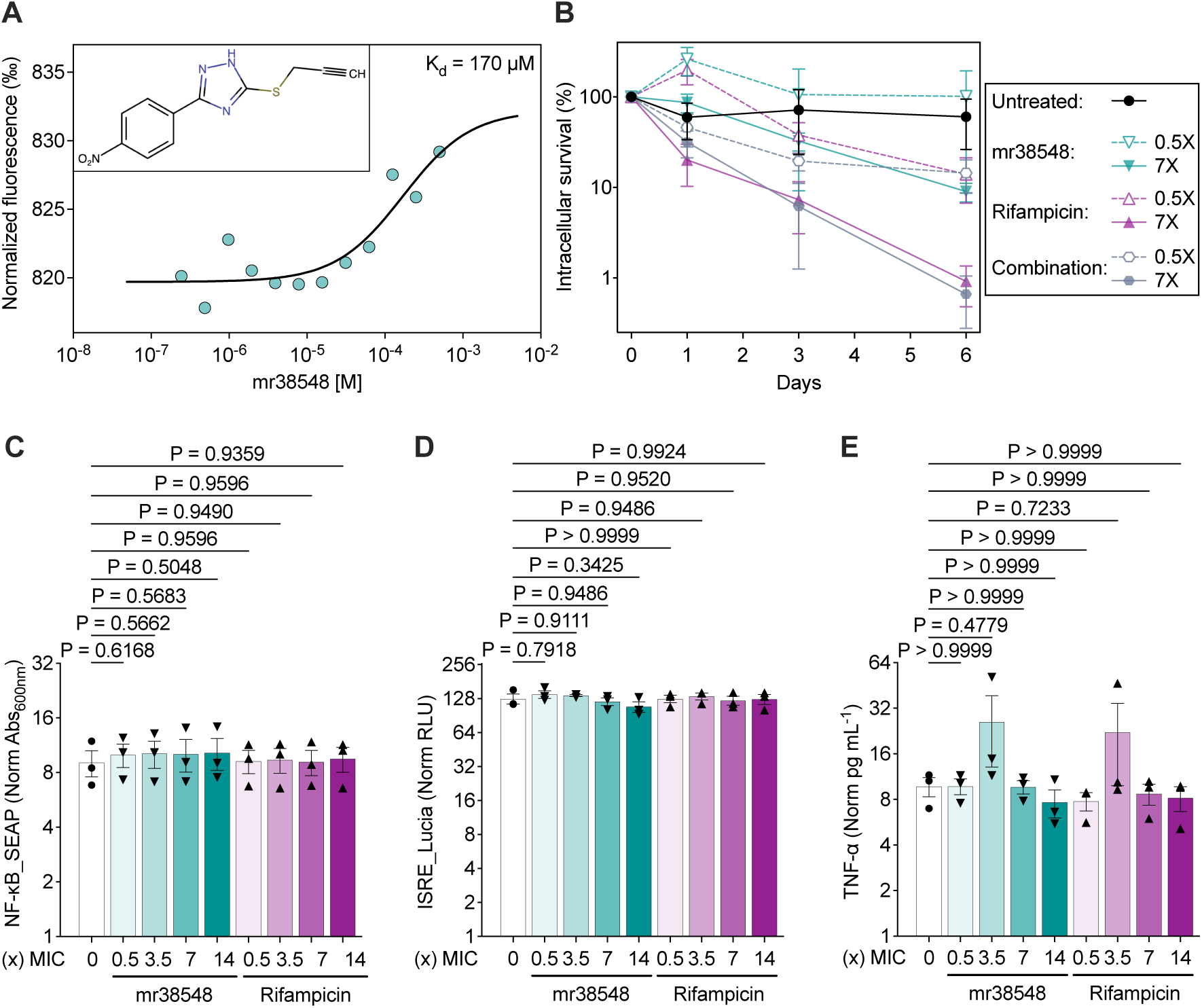
Effect of the putative GroEL2 inhibitor mr38548 at the molecular and cellular levels **(A)** Representative microscale thermophoresis (MST) analysis of binding between GroEL2 binding and mr38548 (n = 3). His-tagged GroEL2 was fluorescently labeled and incubated with increasing concentrations of mr38548 (inset). Labeling was efficient, and no compound autofluorescence, aggregation, or other artifacts were detected. Binding of mr38548 to GroEL2 (80 nM) produced a moderate shift in the MST signal, indicating weak or transient interaction. The dissociation constant (K_d_) is reported on the plot with a confidence interval of 33– 882 µM. **(B)** Intracellular growth of *M. tuberculosis* in THP-1 macrophages infected at MOI 3 or 10. Cells were left untreated (black circles) or treated with mr38548 (green inverted triangles), RIF (purple triangles), or their combinations (gray hexagons) at 0.5x MIC (open symbols and dashed lines) or 7x MIC (filled symbols and solid lines). Data are mean ± SEM (n = 4 independent experiments). **(C–E)** Activation of NF-κB (C) and IRF (D), and TNF-α production (E) in WT THP-1 dual-reporter cells stimulated with LPS (100 ng/mL) for 3 hours. Cells were left untreated (circles) or treated with mr38548 (MIC: 0.88 µg/mL, inverted triangles) or RIF (MIC: 0.076 µg/mL, triangles), at different concentrations relative to their MIC (x). Values are normalized to unstimulated and untreated control cells. Statistics by one-way ANOVA followed by Šidák’s multiple comparisons test (not significant). Lines represent mean ± SEM (n = 3).

## Supplemental video titles and legends

**Video S1. Time-lapse microscopy of *M. tuberculosis* GroEL2_mRuby_-GFP_cyt_ reporter** Representative video of exponential-phase bacteria growing inside our standard microfluidic hexa-device (Manina et al., 2019) and supplied with a constant flow of fresh 7H9 medium at 10 µL/min. Images were recorded at 3-hour intervals (10 fps). Timestamp is shown in hours. Scale bar, 5 μm.

Video S2. Time-lapse microscopy of THP-1 macrophages infected with *M. tuberculosis* GroEL2_mRuby_-GFP_cyt_ reporter

Representative video of resting THP1 macrophages following infection with exponentially growing bacilli at MOI 3 for 2 hours. Infected cells were embedded in a hydrogel layer before assembly with the hexa-device (Manina et al., 2019) and were constantly fed with fresh DMEM medium at 1.8 µL/min. Seven images spanning a z-stack of 12.8 µm were acquired at 4-hours intervals and combined by maximum intensity projection. Timestamp is shown in hours. Scale bar, 10 μm. The field of view shows macrophages infected with either GroEL2_mRuby_-Dim or -Bright bacilli.

## Supplemental table titles and legends

Table S1. Differential gene expression analysis between *M. tuberculosis* FdxA_mCherry_ subpopulations

Raw and normalized gene expression counts (4302 genes) in FdxA_mCherry_-Bright (B) and -Dim (D) subpopulations under each condition tested: exponential phase (EP) and stationary phase (SP) of growth, fatty acids (FA), and hypoxia (HO). Data are from three independent repeats (R). Fold changes of B versus D subpopulations and associated statistics, including Benjamini-Hochberg-corrected P values (padj) are reported for each comparison. The reference condition for each comparison is indicated in the worksheet title. Three worksheets are provided for each condition: full gene set (complete), significant genes upregulated (up) in B, and significant genes downregulated (down) in B.

**Table S2. Functional transcriptome analysis of *M. tuberculosis* FdxA_mCherry_ subpopulations** DEGs in FdxA_mCherry_-Bright (B) versus -Dim (D) subpopulations are reported, either separated or merged by condition: exponential phase (EP), stationary phase (SP), fatty acid (FA), and hypoxia (HO). For each DEG, the *M. tuberculosis* Erdman gene ID and corresponding *M. tuberculosis* H37Rv gene names are provided and classified based on their functional category according to Mycobrowser (Kapopoulou et al., 2011). Functional categories are color-coded as follows: adaptation (light green); lipids (brown); cell wall (dark green); metabolism (orange); regulation (light blue); DNA and RNA (light red); genome insertion (dark red); PE/PPE (purple); conserved (gray); unknown (pink); and unclassified (white). Functionally classified significant DEGs (P < 0.05), with the direction of expression in B versus D subpopulation, are reported for each condition, as well as DEGs upregulated in B that are shared among EP, FA, and HO conditions. Gene ontology (GO) enrichment analyses are provided in separate tabs for DEGs significantly upregulated in B under EP, FA, and HO; for DEG significantly downregulated in B under SP; and for DEG significantly upregulated in B and shared among EP, FA, and HO. Enriched GO biological processes were identified by PANTHER overrepresentation test of significant DEGs against all genes in *M. tuberculosis* H37Rv database (The Gene Ontology Consortium, 2021; Mi et al., 2019). Significance was assessed by Fisher’s exact test followed by Benjamini-Hochberg procedure (FDR < 0.05), and results are sorted by fold enrichment. Bolded GO categories correspond to higher-level, less specific biological processes in the GO hierarchy.

**Table S3.** *M. tuberculosis* nCounter gene expression analysis in patient sputum Raw and normalized gene expression counts are reported for 130 *M. tuberculosis* biomarkers that are differentially upregulated (>2-fold) in the FdxA_mCherry_-Bright versus -Dim subpopulations, shared between at least two conditions, with functions relevant to virulence and stress response, and potentially associated with the cell envelope or secretion. The NanoString probe codeset is provided in the first tab. Data are from ten sputum samples, with 39–99% of probes detected above the negative control threshold on the nCounter Analysis System (NanoString Technologies, Bruker).

Table S4. Mass spectrometry analysis of THP-1 proteins pulled down with GroEL2-His Differentially abundant proteins are separated per condition in THP-1 KO_MyD88_ versus WT cells.

**Table S5.**
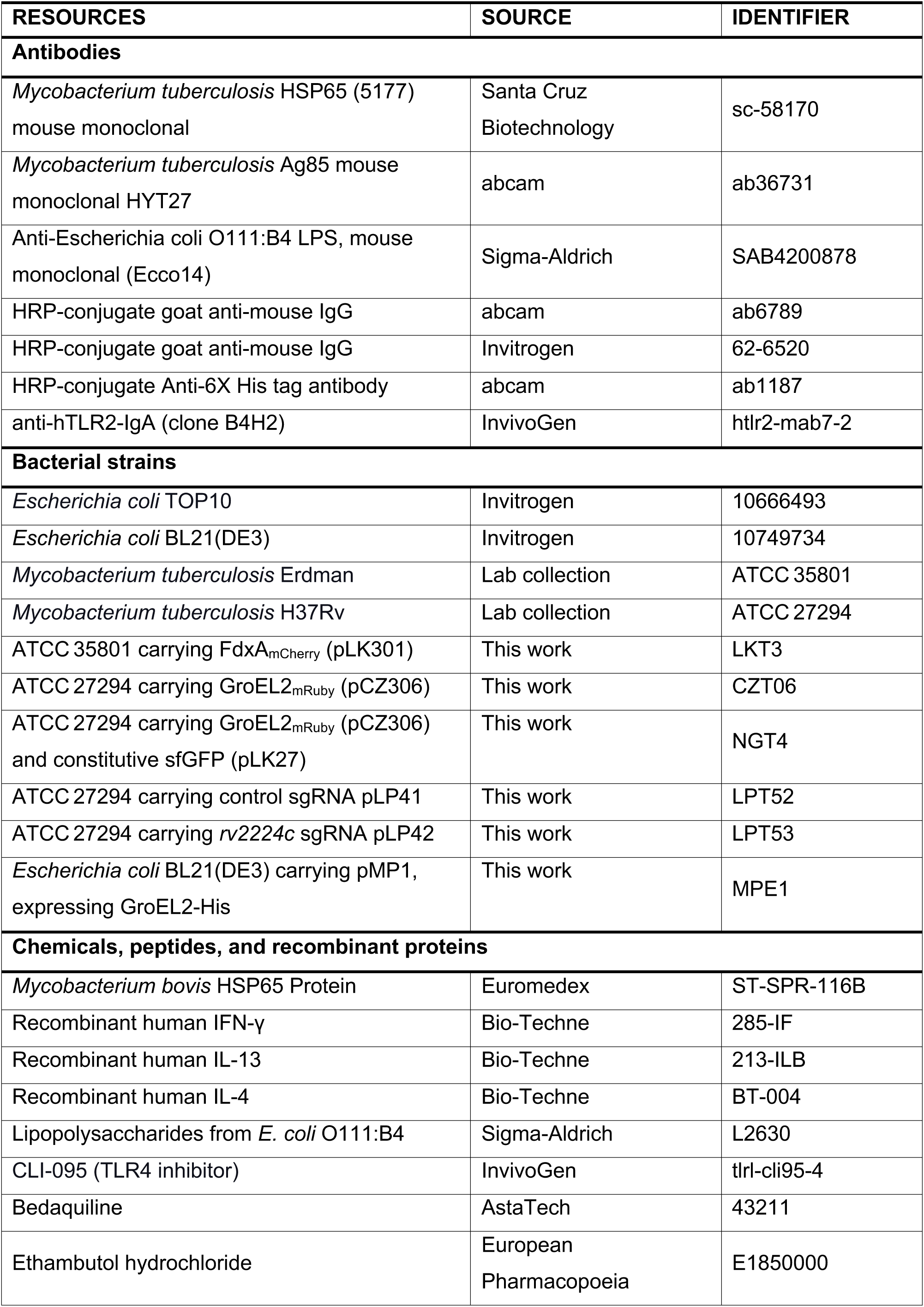
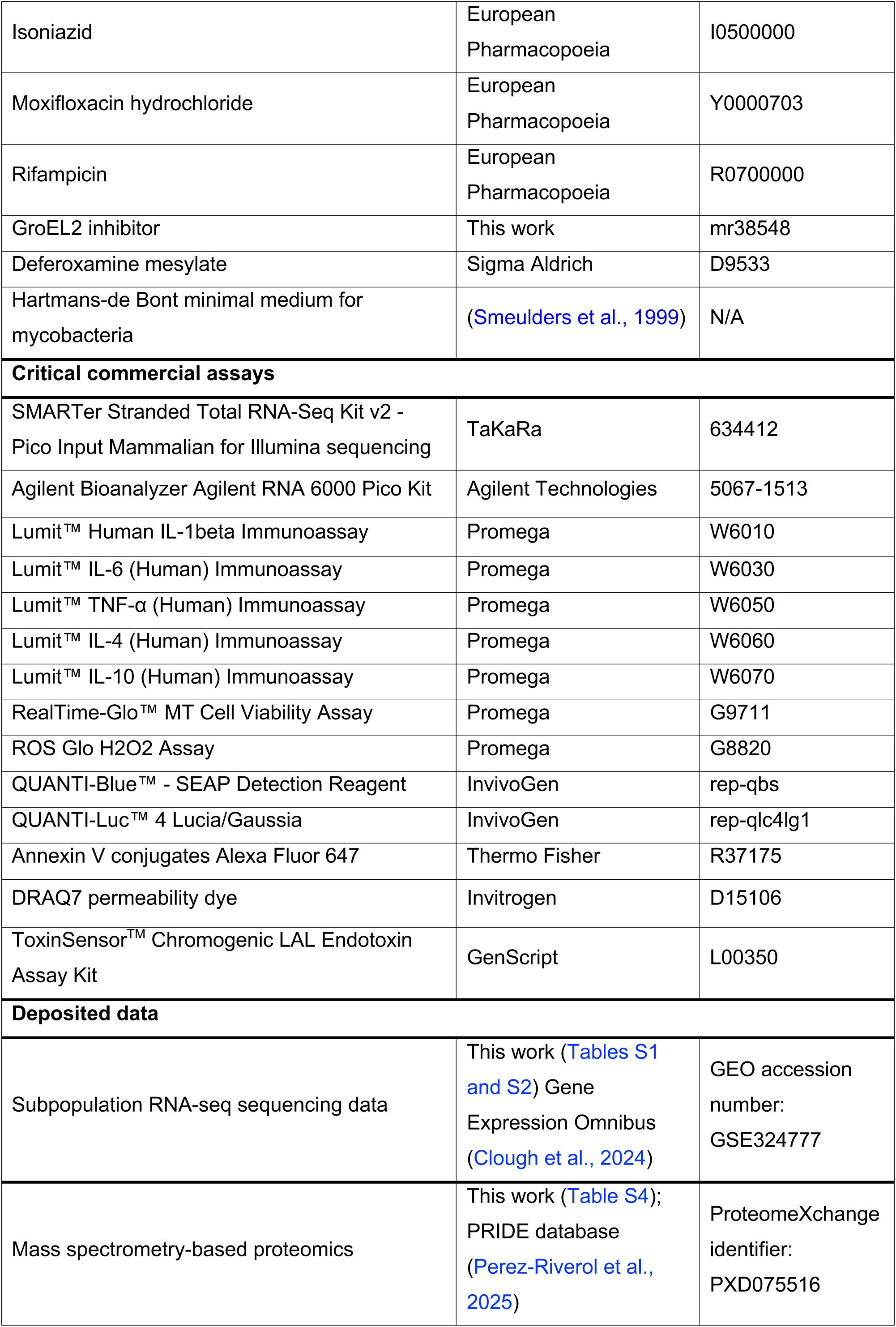
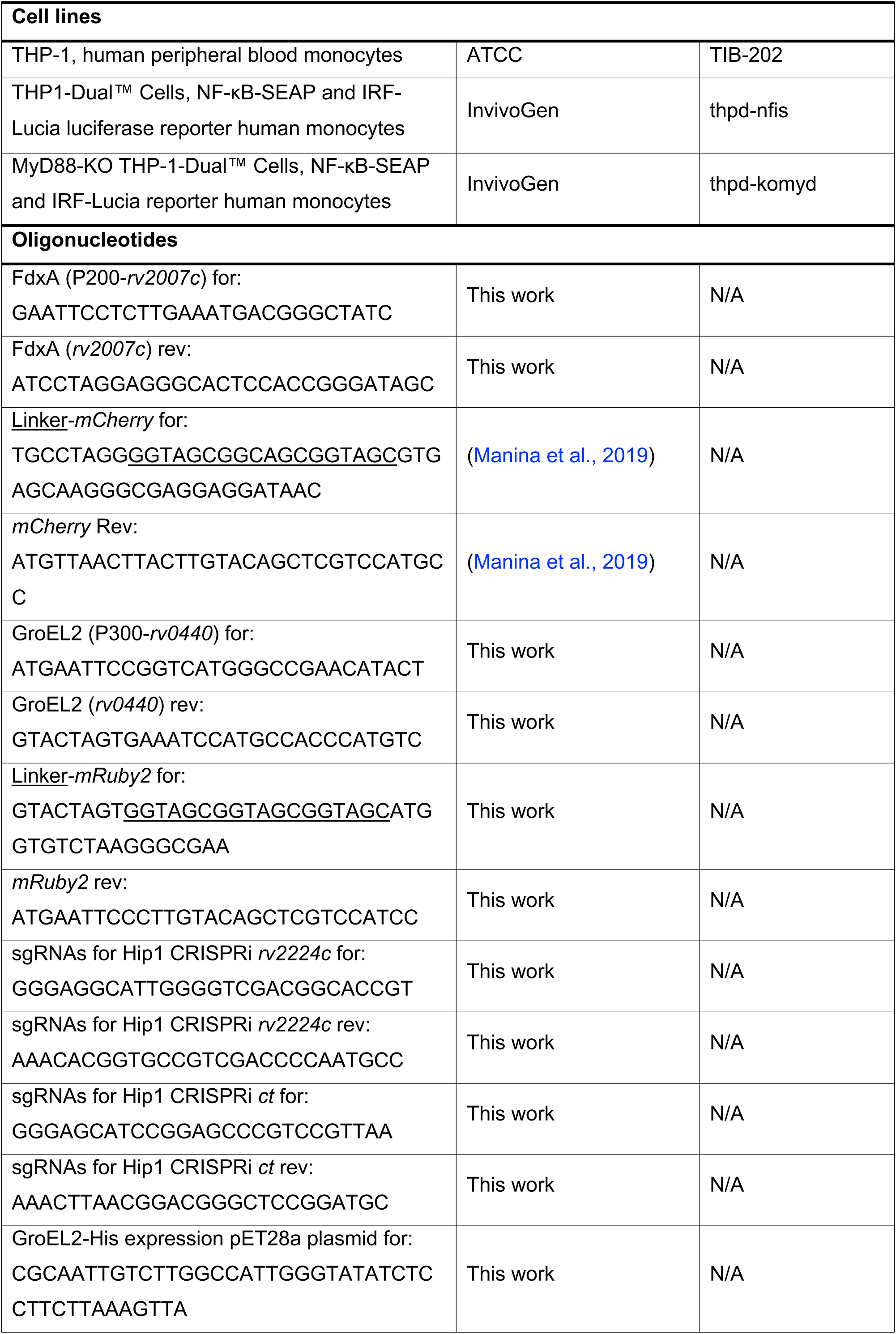
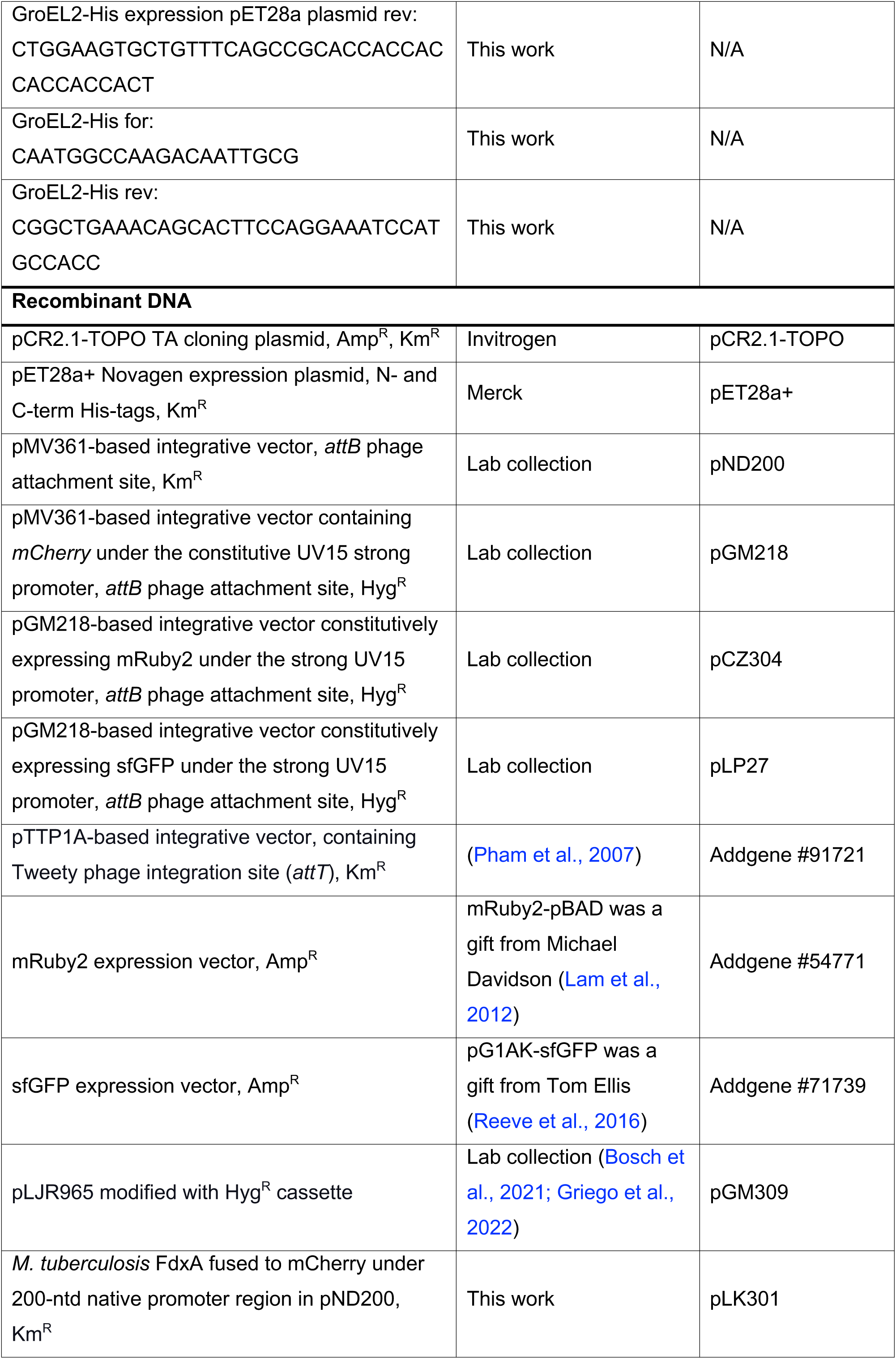
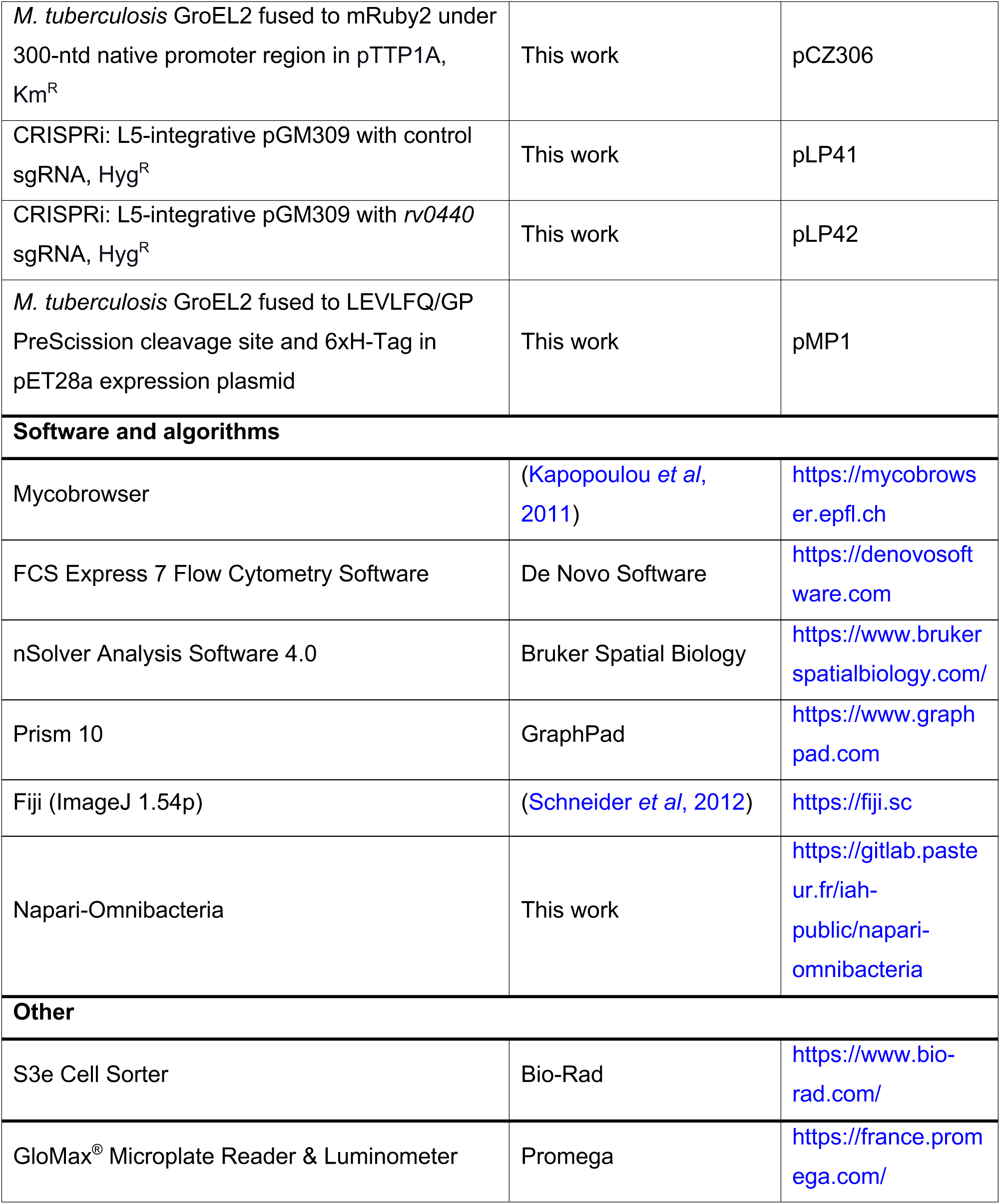
List of reagents and resources used in this study

